# Lysophosphatidic acid-mediated GPR35 signaling in CX3CR1^+^ macrophages regulates the intestinal cytokine milieu

**DOI:** 10.1101/2020.02.03.932186

**Authors:** Berna Kaya, Cristian Doñas Cuadra, Philipp Wuggenig, Oscar E. Diaz, Rodrigo A. Morales, Hassan Melhem, Swiss IBD Cohort Investigators, Pedro P. Hernández, Tanay Kaymak, Srustidhar Das, Petr Hruz, C. Korcan Ayata, Eduardo J. Villablanca, Jan Hendrik Niess

## Abstract

Single nucleotide polymorphisms in the gene encoding G protein-coupled receptor 35 (GPR35) are associated with increased risk of inflammatory bowel disease. However, the mechanism(s) by which GPR35 modulates the intestinal milieu remain undefined. Here we demonstrate in zebrafish and mice that expression of *Gpr35* is microbiota-dependent and is enhanced upon inflammation. We identify a GPR35^+^ colonic macrophage population in mice that is characterized by increased production of pro-inflammatory cytokines, and determine that lysophosphatidic acid (LPA) acts as an endogenous GPR35 ligand to induce *<u>Tnf</u>* expression. Mice lacking *Gpr35* in CX3CR1^+^ macrophages have aggravated colitis when exposed to dextran sodium sulfate, have decreased transcript levels of the corticosterone-generating gene *Cyp11b1*, and reduced levels of macrophage-derived TNF. Administration of TNF in these mice restores *Cyp11b1* expression and intestinal corticosterone production, and ameliorates DSS-induced colitis. These findings suggest that LPA signals through GPR35 in CX3CR1^+^ macrophages to control the intestinal cytokine milieu.

**Highlights:**

1. Inflammatory cues and the microbiota modulate *Gpr35* expression across species
2. LPA modulates GPR35-dependent functions in zebrafish and mice macrophages
3. GPR35 expressing macrophages have a protective role during intestinal inflammation
4. GPR35 control intestinal inflammation by inducing TNF and corticosterone synthesis

**eTOC Blurb:** GPR35 have been associated with IBD, but how GPR35 may influence macrophage-mediated intestinal homeostasis remains unclear. Using zebrafish and mice genetic tools, Niess, Villablanca, and colleagues have identified that LPA triggers GPR35 activity, and loss of macrophage GPR35 signaling confers intrinsic dysfunctions with effects on cytokine production and intestinal homeostasis.

## Introduction

Host- and/or bacterial-derived metabolites orchestrate a wide range of immune responses through G protein-coupled receptors (GPCRs) (Melhem et al., 2019; Postler and Ghosh, 2017; Thorburn et al., 2014). Genome-wide association studies (GWASs) have identified single nucleotide polymorphisms in the coding region of *GPR35* that are associated with increased risk of ulcerative colitis (UC) and primary sclerosing cholangitis (Ellinghaus et al., 2013; Imielinski et al., 2009). Structural modeling studies have suggested that protein-coding variant rs3749171, which lead to the amino acid substitution T108M, may affect the ability of GPR35 to become activated (Ellinghaus et al., 2013); however, how defective GPR35 signaling influence intestinal immune homeostasis is yet poorly understood. The endogenous ligand for GPR35 also remains undefined, although the chemokine CXCL17, the tryptophan metabolite kynurenic acid (KYNA), and the phospholipid derivative lysophosphatidic acid (LPA) have been suggested as putative ligands (Maravillas-Montero et al., 2015; Oka et al., 2010; Wang et al., 2006). Illustrating the complexity of the signaling pathway, some studies have suggested that specific ligands might activate GPR35 in a context- and species-dependent manner. For example, CXCL17 did not activate migration of GPR35-expressing cells in one study (Binti Mohd Amir et al., 2018). Furthermore, KYNA has a wide spectrum of potency for GPR35 across species, with low potency in humans when applied in micromolar concentrations (Mackenzie et al., 2011), and LPA has not been experimentally pursued as a potential GPR35 ligand following the initial suggestion of its role in elevating intracellular Ca(2+) concentrating and inducing receptor internalization (Oka et al., 2010). Thus, the putative GPR35 ligand that might control immune responses in vivo, remain to be identified.

In the intestine GPR35 is highly expressed in the intestinal epithelium and macrophages both in human and mice (Lattin et al., 2008), and activation of GPR35 promotes intestinal epithelial cell turnover during wound healing (Schneditz et al., 2019; Tsukahara et al., 2017). Although these studies used well-known synthetic GPR35 agonists, it remains to be demonstrated if the effect is GPR35 specific both in vitro and in vivo. Kaneider N.C. and colleagues have recently shown that *Gpr35*-deficient compared to control mice resulted in decreased inflammation-associated intestinal tumorigenesis (Schneditz et al., 2019; Tsukahara et al., 2017). In addition, Gpr35^-/-^ intestinal epithelial cells resulted in reduced turnover compared to control mice, as seen by proliferation analysis in situ and in organoids cultures (Schneditz et al., 2019; Tsukahara et al., 2017). Thus, indicating the GPR35 might orchestrate homeostatic epithelial cell renewal. Moreover, in agreement with a role in IEC turnover, GPR35 is protective during chemically induced acute colonic inflammation in mice (Farooq et al., 2018). However, the endogenous ligand and cell type triggering GPR35 signaling to maintain intestinal homeostasis remains unknown.

Several lines of evidence have implicated macrophages in inflammatory bowel disease (IBD). For example, multiple IBD risk genes play critical roles in macrophages functions, such as bacterial clearance (e.g. *Gpr65* and *Nod2*) (Hedl and Abraham, 2011; Lassen et al., 2016; Peters et al., 2017). Importantly, the critical role of macrophages in the establishment of intestinal homeostasis have been demonstrated by targeted depletion of tolerogenic signals in macrophages which results in spontaneous colitis in mice (Bernshtein et al., 2019; Shouval et al., 2014; Zigmond et al., 2014). Macrophages are also part of the inflammatory cell infiltrates in mice with colitis and in patients with IBD, suggesting that macrophages might not only prevent but also drive intestinal inflammation by producing pro-inflammatory cytokines, such as IL-6, IL-1β, and TNF (MacDonald et al., 2011). GPR35 is also expressed in bone marrow-derived macrophages (BMDMs), and peritoneal macrophages (Lattin et al., 2008; Schneditz et al., 2019; Tsukahara et al., 2017). GPR35 interact and modulate the activity of Na/K-ATPase to eventually control macrophage metabolism (Schneditz et al., 2019; Tsukahara et al., 2017). However, if GPR35 signaling in macrophages is critical to establish intestinal immune homeostasis in vivo and the putative ligand that may trigger GPR35-dependent functions in macrophages is yet to be understood.

TNF has drawn special attention in the study of IBD since TNF can drive colitis and targeting TNF with antibodies attenuates IBD in the clinic. For instance, inhibition of TNF prevents the progression of symptoms in several mouse models of disease, including spontaneous ileitis (Gunther et al., 2011), 2,4,6-trinitrobenzenesulfonic acid (TNBS) colitis (Neurath et al., 1997), and transfer colitis (Corazza et al., 1999). Anti-TNF antibodies have also been successfully implemented in the clinic for the treatment of patients with IBD (Hanauer et al., 2002; Targan et al., 1997). Besides the well-described pro-inflammatory role of TNF in colitis, some evidence suggests that TNF also has anti-inflammatory functions. For example, in the DSS colitis model, the neutralization of TNF or absence of TNF in *Tnf*-deficient mice leads to an exacerbation of colitis (Naito et al., 2003), an effect that has been attributed to TNF-mediated induction of apoptosis in T cells, which leads to resolution of inflammation (Zheng et al., 1995). TNF has also been suggested to regulate extra-adrenal corticosterone production by intestinal epithelial cells, which in turn may suppress immune responses (Noti et al., 2010).

Here we report that high *Gpr35* expression in the gastrointestinal tract is conserved across species and that GPR35 distinguishes two intestinal lamina propria CX3CR1^+^ macrophage subpopulations that are transcriptionally distinct. Using in vitro and in vivo approaches in *gpr35* mutant zebrafish, *Gpr35*-deficient mice, and cells expressing human GPR35, we identify LPA as an endogenous ligand that triggers GPR35-dependent induction of *Tnf* and *Il1b* transcripts in macrophages. Conditional deletion of *Gpr35* in macrophages in mice results increased susceptibility to DSS-induced colitis, reduced TNF production by CX3CR1^+^ macrophages in vivo, and decreased *Cyp11b1* expression. Finally, TNF administration into mice lacking *Gpr35* in macrophages attenuated DSS-induced colitis and restored *Cyp11b1* expression. Thus, our data indicate that GPR35 controls macrophage function to maintain the intestinal cytokine milieu under steady-state conditions and during intestinal inflammation.

## Results

### Colonic Macrophages Express GPR35

Our interest in using zebrafish (*Danio rerio*) to investigate the function of IBD-associated risk genes led us to clone and characterize the functional zebrafish homolog of *GPR35*. We identified two *GPR35* paralogs in zebrafish that we named *gpr35a* and *gpr35b*, which share 25.6% and 24% identity with the human GPR35 protein sequence, respectively (Figure S1A). Of note, *gpr35a* and *gpr35b* were more closely related phylogenetically to human *GPR35* and murine *Gpr35* than to human or murine versions of *GPR55*, a highly similar gene (Figure S1B). Gene expression analysis revealed that *gpr35a* was expressed at similar levels in the intestine compared to the rest of the body, whereas *gpr35b* was predominantly expressed in the intestine by 120 hours post fertilization (hpf) (Figures 1A and S1C). This finding was confirmed by whole in situ hybridization (WISH), which showed expression of *gpr35b* specifically in the intestinal bulb at 120 hpf (Figure 1B). These observations were echoed in our analysis of the human protein atlas, which showed the highest *GPR35* transcript levels in the gastrointestinal tract compared to other tissues (Figure S1D). Similarly, qRT-PCR of mouse tissues revealed an increased expression pattern of *Gpr35* from the duodenum to the distal colon, in the proximal stomach, mesenteric lymph nodes (MLN), and Peyer’s patches, compared to other tissues, such as the liver (Figure 1C).

**Figure 1.**
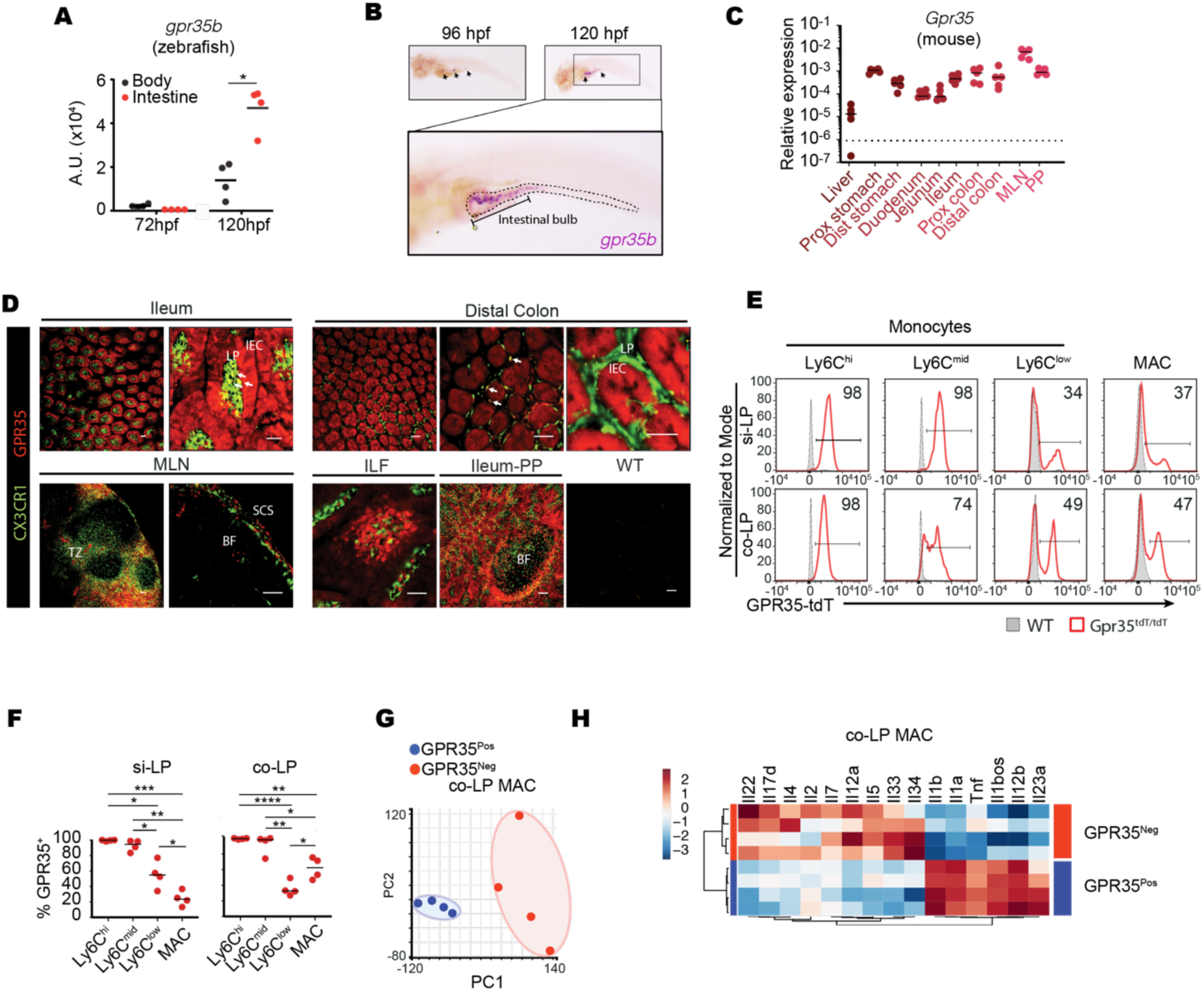
GPR35 Is Expressed in Colonic Macrophages. (A) *gpr35b* mRNA expression levels by qRT-PCR normalized to *ef1a* across the body and dissected intestines of zebrafish larvae at 72 hpf and 120 hpf. A.U.; arbitrary units normalized to the lower value (body, 72 hpf). (B) Whole mount in situ hybridization (WISH) to detect *gpr35b* mRNA expression in zebrafish larvae at 96 hpf and 120 hpf. Arrows indicate the intestinal bulb; dashed lines indicate the intestinal tract. One representative picture is shown from 40 larvae. (C) *Gpr35* mRNA expression levels by qRT-PCR normalized to *Gapdh* across indicated tissues in WT mice. (D) Ex vivo fluorescence imaging of ileum, distal colon, mesenteric lymph node (MLN), isolated lymph follicle (ILF), and Peyer’s patch (PP) from *Cx3cr1*-GFP (green) x *Gpr35*-tdTomato (red) double reporter mice. The last panel shows colon from WT as the background control. LP, lamina propria; IEC, intestinal epithelial cell; TZ, T cell zone; BF, B cell follicle; SCS, Subcapsular sinus. Arrows indicate CX3CR1^+^ phagocytes that express GPR35. Scale bars, 50 μm. (E) Representative *Gpr35*-tdTomato expression by flow cytometry in monocyte subsets (Ly6C^high^ to Ly6C^low^) and macrophages (MAC) from small intestinal and colonic lamina propria (si-LP and co-LP) of *Gpr35*-tdTomato reporter mice (red unfilled histograms) and WT mice (gray histograms) as the background control. Numbers in histograms indicate the percentage of GPR35-tdTomato^+^ cells. (F) Quantification of data from (E) showing the percentage of GPR35-tdTomato^+^ cells in monocytes and macrophages in the si-LP and co-LP. (G) Principal component analysis from RNA sequencing of GPR35-tdTomato-positive (GPR35^pos^) and -negative (GPR35^neg^) colon lamina propria (co-LP) macrophages. (H) Heatmap representation of cytokine expression profiles from RNA sequencing of GPR35-tdTomato-positive (pos) and -negative (neg) subpopulations in co-LP macrophages. Data are represented as individual values with medians with each dot representing one biological replicate. *p ≤ 0.05, **p ≤ 0.01, ***p ≤ 0.001, ****p ≤ 0.0001 by two-way ANOVA with Tukey’s multiple comparisons test.

To more precisely define the population of cells expressing GPR35, we generated a *Gpr35*-tdTomato reporter mouse line (Figure S2A). Immunofluorescent staining for GPR35 showed colocalization with td-Tomato, indicating that the reporter accurately monitored endogenous *Gpr35* expression (Figure S2B). Ex vivo imaging of small and large intestinal tissues from *Gpr35*-tdTomato mice crossed to *Cx3cr1*-GFP reporter mice revealed GPR35 expression in intestinal epithelial cells and lamina propria CX3CR1^+^ phagocytes (Figure 1D). In addition, GPR35^+^ cells were located in subcapsular sinus and T cell zones of MLN, in isolated lymph follicles and the subepithelial dome regions of Peyer’s patches, but not in B cell follicles of MLN neither/nor in Peyer’s patches (Figure 1D). We also observed GPR35 expression in CD64^-^CD11c^+^ dendritic cells, whereas no expression was detected in B cells, CD4^+^ or CD8^+^ T cells, neutrophils, NK cells, or innate lymphoid cells, including ILC1, ILC2, and ILC3 cells (Figure S2C).

To further track the expression of GPR35 in macrophages and their precursors, we analyzed the expression of GPR35 by macrophages and monocytes with flow cytometry. CX3CR1^+^ macrophages derive from blood Ly6C^high^ monocytes that extravasate into the lamina propria, downregulate Ly6c and develop into mature macrophages through intermediates in in a “monocyte waterfall” development (Bain et al., 2013). When we examined the expression of GPR35 alongside the differentiation of monocytes into macrophages (Steinert et al., 2017) (Figure S2D), most Ly6C^high^ monocytes in the lamina propria of the small and large intestine expressed GPR35 (Figure 1E). Percentages of GPR35^+^ cells gradually decreased to approximately 30% and 50% alongside the differentiation of monocytes into mature macrophages in the small and large intestine, respectively (Figure 1F) suggesting that monocytes down-regulate GPR35 during their maturation into colonic macrophages.

To gain insight into the potential function of GPR35^+^ macrophages, we performed bulk RNA-seq analysis on sorted GPR35^-^ and GPR35^+^ macrophages from the colonic lamina propria. Unsupervised hierarchical clustering and principal component analysis (PCA) revealed that GPR35^-^ and GPR35^+^ macrophages are transcriptionally distinct populations (Figures 1G and S2E), with GPR35^+^ macrophages showing higher *Il1b*, *Il1a*, *Tnf*, *Il12b*, and *Il23a* transcript levels compared to GPR35^-^ macrophages (Figure 1H). Taken together, these data show that GPR35 is highly expressed in intestinal tissues across species, and GPR35^+^ murine colonic macrophages are characterized by higher expression of pro-inflammatory genes compared to GPR35^-^ macrophages.

### *Gpr35* Expression Is Microbiota-Dependent and Upregulated Upon Inflammation

Next, we considered the possibility that the intestinal environment could modulate the expression of GPR35. To address this question, we first investigated whether induction of *gpr35* mRNA expression in the zebrafish intestine was microbiota-dependent. We found that perturbation of intestinal bacteria by treatment with antibiotics resulted in reduced intestinal *gpr35b* transcript levels, as seen by WISH and qPCR, when compared to vehicle-treated zebrafish (Figures 2A and 2B). Analogous results were obtained in mice, in which administration of a broad-spectrum antibiotic cocktail (vancomycin, neomycin, metronidazole, gentamicin, and ampicillin) resulted in decreased levels of *Gpr35* transcript in the colonic lamina propria compared to non-treated mice (Figure 2C). Similarly, colonic tissue from germ-free mice had lower *Gpr35* expression levels compared to specific pathogen-free mice (Figure 2D). Altogether, these results suggest that the microbiota modulates intestinal *Gpr35* expression in zebrafish and mice.

**Figure 2.**
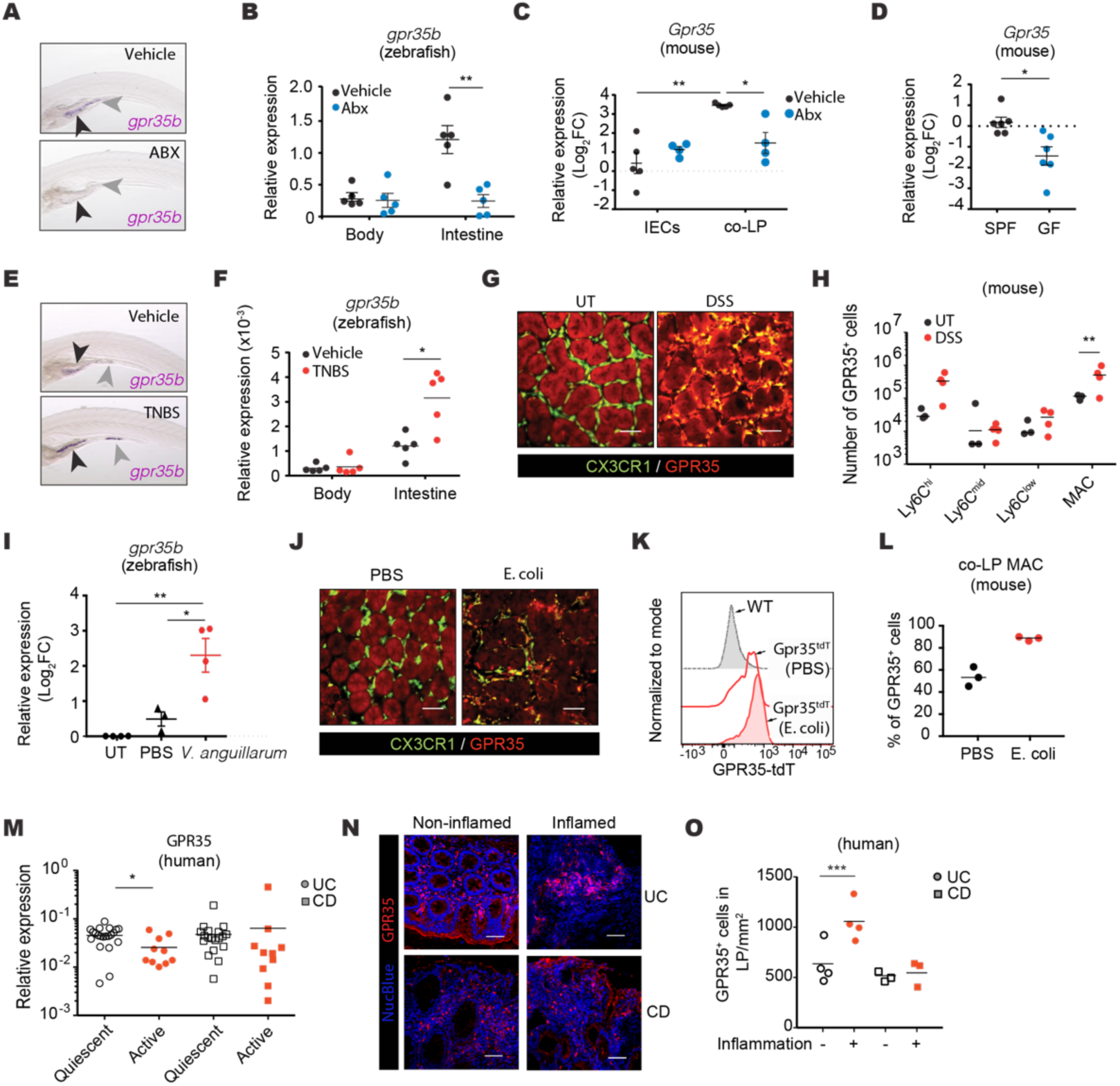
*Gpr35* Expression is Modulated by the Microbiota and Inflammation. (A) *gpr35b* mRNA expression detected by WISH using a *gpr35b* anti-sense probe in 120 hpf zebrafish larvae treated with antibiotics (ABX) or vehicle. Larvae were treated for 48 h starting at 72 hpf; antibiotics were diluted in E3 water. Arrowheads indicate the intestinal bulb. One representative picture is shown from 40 larvae. (B) *gpr35* mRNA expression levels by qRT-PCR normalized to *ef1a* across the body and dissected intestines from WT zebrafish treated with antibiotics or vehicle as described in (A). Each dot represents a pool of 10 larvae. (C) *Gpr35* mRNA expression levels by qRT-PCR normalized to *Hprt* in intestinal epithelial cells (IECs) and colonic lamina propria cells (co-LP) from WT mice treated with an antibiotics cocktail or vehicle. Antibiotics were administered every 24 h for 10 days by oral gavage. (D) *Gpr35* mRNA expression levels by qRT-PCR normalized to *Hprt* in colonic tissue from specific pathogen-free (SPF) and germ-free (GF) mice. (E) *gpr35b* mRNA expression detected by WISH in 120 hpf zebrafish larvae treated with TNBS or vehicle. Larvae were treated for 48 h starting at 72 hpf; TNBS was diluted in E3 water. Arrowheads indicate intestinal bulb and the posterior intestine. One representative picture is shown from 20 larvae. (F) *gpr35b* mRNA expression levels by qRT-PCR normalized to *ef1a* across the body and dissected intestines from WT zebrafish treated with TNBS or vehicle as described in (E). Each dot represents a pool of 10 larvae. (G) Ex vivo fluorescence imaging of colon from untreated (UT) or DSS-treated *Cx3cr1*-GFP (green) x *Gpr35*-tdTomato (red) double reporter mice on day 7 of DSS colitis. Scale bars, 50 μm. (H) Numbers of GPR35-tdTomato^+^ monocytes (Ly6C^high^ to Ly6C^low^) and macrophages (MAC) quantified by flow cytometry of co-LP from UT and DSS-treated *Gpr35*-tdTomato mice. (I) *gpr35b* mRNA expression levels measured by qRT-PCR normalized to *ef1a* in WT zebrafish exposed to PBS or *V. anguillarum*. Injections of *V. anguillarum* extracts were performed in the swim bladder/intestine of 112 hpf anesthetized larvae; tissues were harvested 6 h post injection to isolate total mRNA. Each dot represents a pool of 10 larvae. (J) Ex vivo fluorescence imaging of colon from *Cx3cr1*-GFP (green) x *Gpr35*-tdTomato (red) mice gavaged with PBS or *E. coli* every other day; tissue collected on day 21. Scale bar, 50 μm. (K) Representative *Gpr35*-tdTomato expression by flow cytometry in co-LP macrophages from WT (gray histogram), PBS-gavaged *Cx3cr1*-GFP x *Gpr35*-tdTomato (red unfilled histogram), and *E. coli*-gavaged *Cx3cr1*-GFP x *Gpr35*-tdTomato (red filled histogram) mice. (L) Quantification of flow cytometric data from (K) for numbers of GPR35^+^ cells in co-LP macrophages in PBS-gavaged or *E. coli*-gavaged *Cx3cr1*-GFP x *Gpr35*-tdTomato mice. (M) *GPR35* mRNA expression by qRT-PCR comparing biopsies from ulcerative colitis (UC) or Crohn’s disease (CD) patients with quiescent or active disease. (N) Immunofluorescence imaging of UC or CD patient biopsies taken from non-inflamed (left) or inflamed regions (right). Sections were stained for GPR35 (red) and NucBlue (blue) for nuclear staining. Scale bars, 50 μm. (O) Number of GPR35^+^ cells in the lamina propria per mm^2^ quantified by manual counting of immunofluorescence images as shown in (N) of non-inflamed and inflamed UC or CD biopsies. Data are represented as individual values with medians with each dot representing one biological replicate. *p ≤ 0.05, **p ≤ 0.01, ***p ≤ 0.001, ****p ≤ 0.0001 by unpaired t-test (D), one-way (I) or two-way ANOVA with Tukey’s multiple comparisons test (B, C, F, H, O) or Mann-Whitney (L, M).

Given that the intestinal epithelium and immune system are in constant exposure to inflammatory stimuli from the luminal content and Gpr35 is highly expressed in the intestine, we next hypothesized that *Gpr35* expression might be modulated by inflammation. Supporting this hypothesis, we found that triggering intestinal inflammation by treating zebrafish with TNBS resulted in increased *gpr35b* transcript levels in the intestinal bulb (Figure 2E, black arrowheads) and ectopic expression in the posterior intestine (Figure 2E, grey arrowheads) as observed by WISH. Furthermore, qPCR revealed higher levels of *gpr35b* transcripts in the intestine of TNBS-treated zebrafish compared to vehicle-treated animals (Figure 2F). Similarly, ex vivo imaging of *Cx3cr1*-GFP x *Gpr35-*tdTomato double reporter mice revealed increased *Gpr35*-tdTomato signal by CX3CR1^+^ mononuclear phagocytes in the colon of mice treated with DSS (Figure 2G), and flow cytometric analysis confirmed increased number of GPR35^+^ colonic macrophages in response to DSS (Figure 2H). To determine whether infection-induced colitis would affect *Gpr35* expression similarly to chemically (i.e., DSS) induced colitis, we next injected swim bladders from zebrafish with *Vibrio anguillarum* extracts. As we predicted, zebrafish injected with *V. anguillarum* showed a ∼4-fold increase in *gpr35b* transcripts compared to PBS-treated fish (Figure 2I). In mice, we found that colonization of *Cx3cr1*-GFP x *Gpr35-*tdTomato double reporter mice with *Escherichia coli* DH10B pCFP-OVA (Rossini et al., 2014) induced GPR35 expression in colonic lamina propria macrophages (Figures 2J-2L), providing further evidence that GPR35 expression is modulated in the context of inflammation.

To pursue these results, we next sought to test the clinical relevance of GPR35 upregulation during human colitis. For these studies, we determined the expression of *GPR35* in biopsies provided by the Swiss IBD Cohort Study Group obtained from patients with active or quiescent Crohn’s disease or ulcerative colitis. We found decreased *GPR35* expression in patients with ulcerative colitis with active disease compared to those with quiescent disease; in contrast, patients with Crohn’s disease showed comparable *GPR35* expression between active and quiescent disease in entire tissues (Figure 2M). Since both intestinal epithelial cells and macrophages express GPR35, we next performed staining for GPR35 in patient biopsies to exclude the possibility that *GPR35* expression might be differentially regulated in distinct compartments. In these comparisons, we determined the number of GPR35^+^ cells in the lamina propria of biopsies taken from inflamed or non-inflamed regions of the intestine from the same patients. We found no differences in the numbers of GPR35^+^ cells between inflamed and non-inflamed regions in patients with Crohn’s disease, but we did observe an increase in the numbers of GPR35^+^ cells in inflamed regions compared to non-inflamed segments of the same patient with ulcerative colitis (Figures 2N and 2O). Taken together, these data indicated that lamina propria cells of patients with ulcerative colitis with active disease increase GPR35 expression.

### LPA Induces *Tnf* Expression in Macrophages in a GPR35-Dependent Manner

To identify endogenous ligands of GPR35 in the context of intestinal immunity, we began by focusing on LPA, KYNA, and CXL17, which have been previously suggested as GPR35 ligands (Maravillas-Montero et al., 2015; Oka et al., 2010; Wang et al., 2006). We first screened the activating potential of these candidate ligands using a Chinese hamster ovary (CHO)-K1 GPR35 Gi cell line in which human GPR35 is stably overexpressed and naturally coupled to an inhibitory G protein that inhibits forskolin-induced cAMP accumulation in response to GPR35 agonists. As expected, stimulation with the synthetic GPR35 agonist zaprinast inhibited forskolin-induced cAMP production (Figure 3A). KYNA did not elicit a significant response, whereas LPA and CXCL17 inhibited cAMP production, with LPA exerting its effect at a lower concentration compared to CXCL17 (Figure 3A). We next took advantage of CRISPR/Cas9-based genome engineering of zebrafish (Li et al., 2016) to generate a *gpr35b* mutant line, named *gpr35b^uu1892^* (Figures S3A and S3B). Forty-eight hours of LPA treatment resulted in elevated expression of pro-inflammatory cytokines, including *tnf*, *il1b*, and *il17a/f* in WT fish; however, none of these cytokines were induced by LPA in the *gpr35b^uu19b2^* mutants (Figure 3B), indicating that the LPA-induced expression of cytokines is *gpr35b*-dependent. To validate these findings in mice, we used CRISPR/Cas9 to generate a *Gpr35* knockout (KO) mouse line (Figure S4A) that failed to show GPR35 staining by immunofluorescence (Figure S4B). In line with the zebrafish data, stimulation of WT murine BMDMs with LPA significantly induced *Tnf, Il1b*, and *Il23a* expression, whereas LPA stimulation in *Gpr35*^-/-^ BMDMs did not significantly induce the expression of these cytokines compared to unstimulated *Gpr35*^-/-^ BMDMs (Figure 3C). In addition, we observed a significant difference in *Tnf* transcript levels between WT and G*pr35*^-/-^ BMDMs stimulated with LPA (Figure 3C), suggesting that LPA-mediated *Tnf* induction is dependent on GPR35 expression.

**Figure 3.**
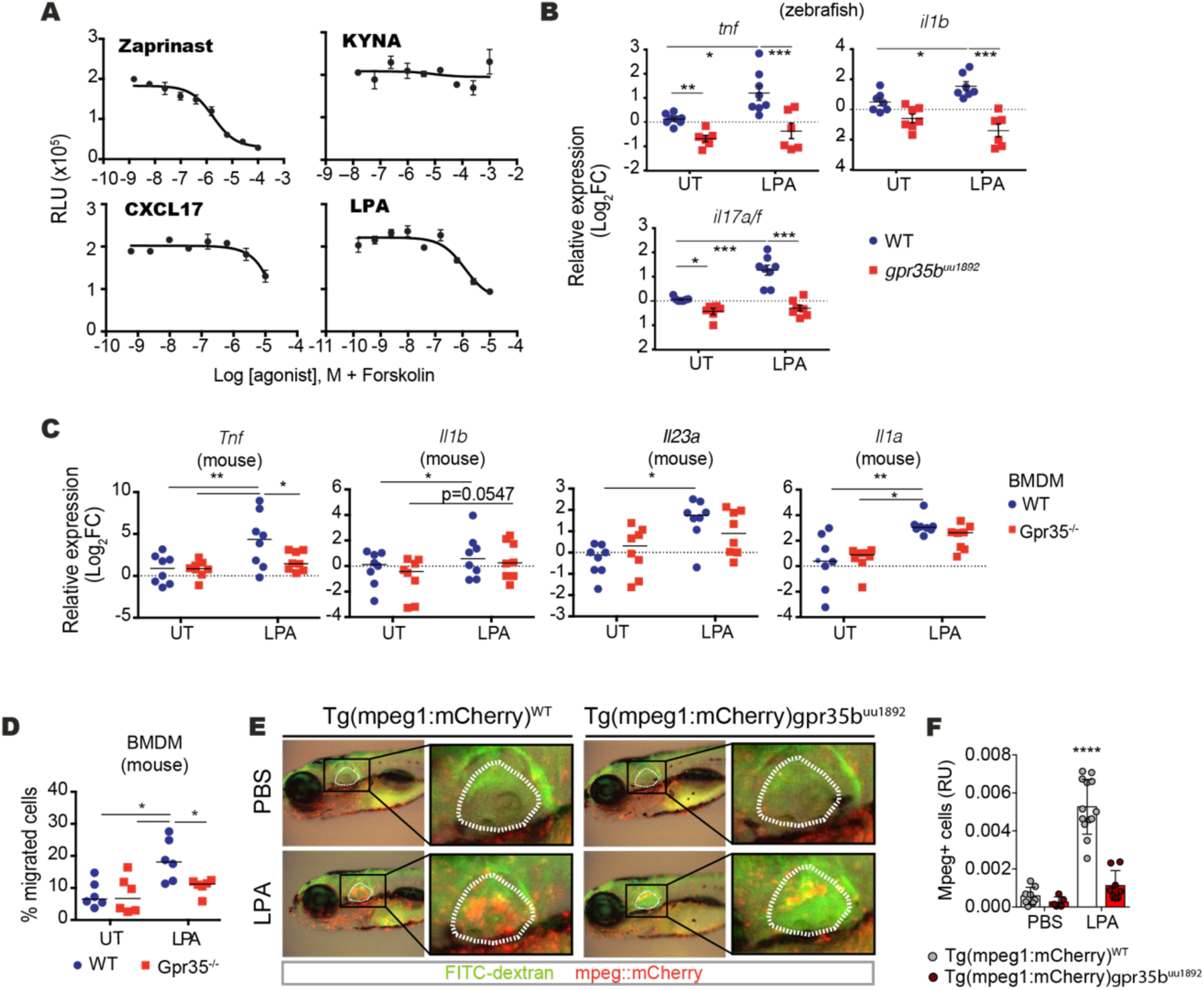
LPA Induces *Tnf* Expression in a GPR35-Dependent Manner. (A) Relative luminescence unit (RLU) values for intracellular cAMP levels in Gi-coupled GPR35-transfected CHO-K1 cells in response to adenylyl cyclase activating-forskolin against serial dilutions of zaprinast, KYNA, CXCL17, or LPA. Data are represented as median ± range from doublets with nonlinear fit curves. (B) mRNA expression levels of *tnf*, *il1b*, and *il17a/f* measured by qRT-PCR in untreated (UT) or LPA-treated WT or *gpr35b^uu1892^* zebrafish larvae. Larvae were treated for 48 h starting at 72 hpf. Analysis was performed at 120 hpf and each dot represents a pool of 10 larvae. (C) mRNA expression levels of *Tnf*, *Il1b*, *Il23a*, and *Il1a* measured by qRT-PCR in UT or LPA-treated bone marrow-derived macrophages (BMDMs) derived from WT or *Gpr35*^-/-^ mice. Results are cumulative of three independent experiments in which each dot represents one mouse. (D) Percentages of migrated BMDMs from WT or *Gpr35*^-/-^ mice towards UT control or LPA in transwell assay acquired by flow cytometry. Results are cumulative of three independent experiments in which each dot represents one mouse. (E) Macrophage recruitment (mpeg1+ cells) in *Tg(mpeg1:mCherry)^WT^* and *Tg(mpeg1:mCherry)gpr35b^uu1892^* zebrafish larvae injected with DMSO or LPA (10 μM) in the otic vesicle (white dashed line). Results are cumulative of two independent experiments in which every dot represents one embryo. (F) Quantification of macrophages recruitment data as shown in (E) for numbers of mpeg1^+^ cells in otic vesicles of *Tg(mpeg1:mCherry)*-WT and *Tg(mpeg1:mCherry)*-*gpr35b^uu1892^* zebrafish larvae injected with PBS or LPA. Data are represented as individual values with medians. *p ≤ 0.05, **p ≤ 0.01, ***p ≤ 0.001, ****p ≤ 0.0001 by two-way ANOVA with Tukey’s multiple comparisons test.

Given that LPA has been previously shown to cause increased migration of monocytes, microglia, and ovarian cancer cells (Oh et al., 2017; Plastira et al., 2017; Takeda et al., 2019), we next tested whether LPA acts as a chemoattractant for macrophages by quantifying the migration of WT and *Gpr35* KO BMDMs in response to LPA in vitro. These experiments revealed that *Gpr35*-deficient BMDMs had reduced migration in response to LPA as compared to WT BMDMs (Figure 3D). To investigate the LPA-GPR35 axis in modulating macrophage chemotaxis in vivo, we next crossed *gpr35b^uu1892^* mutant zebrafish with the reporter strain *Tg(mpeg1:mCherry)* to visualize macrophage dynamics as previously described (Nguyen-Chi et al., 2015). Injection of LPA within the otic vesicle resulted in increased macrophage infiltration compared to PBS injection in WT reporter fish. By contrast, macrophages in *gpr35b^uu1892^* mutant fish did not respond to LPA injection (Figures 3E and 3F), indicating that LPA induces chemotaxis of macrophages in vivo in a Gpr35-dependent fashion.

### Intestinal Inflammation Increases Autotaxin Expression in Zebrafish and Mice

LPA is a phospholipid derivate found in cell membranes and cell walls that can act as an extracellular signaling molecule (Ye and Chun, 2010). LPA is mainly synthesized by autotaxin (ATX), which removes a choline group from lysophosphatidylcholine (Gesta et al., 2002). We therefore investigated whether ATX is induced during intestinal inflammation in vivo using a TNBS model of colitis in zebrafish, which revealed a two-fold increase in *atx* transcripts in intestinal tissues isolated from TNBS-treated zebrafish larvae compared to untreated larvae (Figure 4A). Pursuing these findings in mice, we first consulted our published longitudinal transcriptomic data from mice undergoing DSS colitis (Czarnewski et al., 2019), which showed transient colonic *Atx* expression peaking at d10 (Figure 4B). Consistent with these observations, mice with DSS-induced intestinal inflammation showed an increased number of ATX^+^ cells in colonic tissues as inflammation progressed (Figures 4C and 4D). Notably, *Gpr35^-/-^* mice exposed to DSS showed a comparable increase in the number of ATX^+^ cells compared to WT mice exposed to DSS (Figure 4E), suggesting that GPR35 does not modulate ATX expression. In conclusion, colitis in zebrafish and mice led to increased *Atx* expression in the colon.

**Figure 4.**
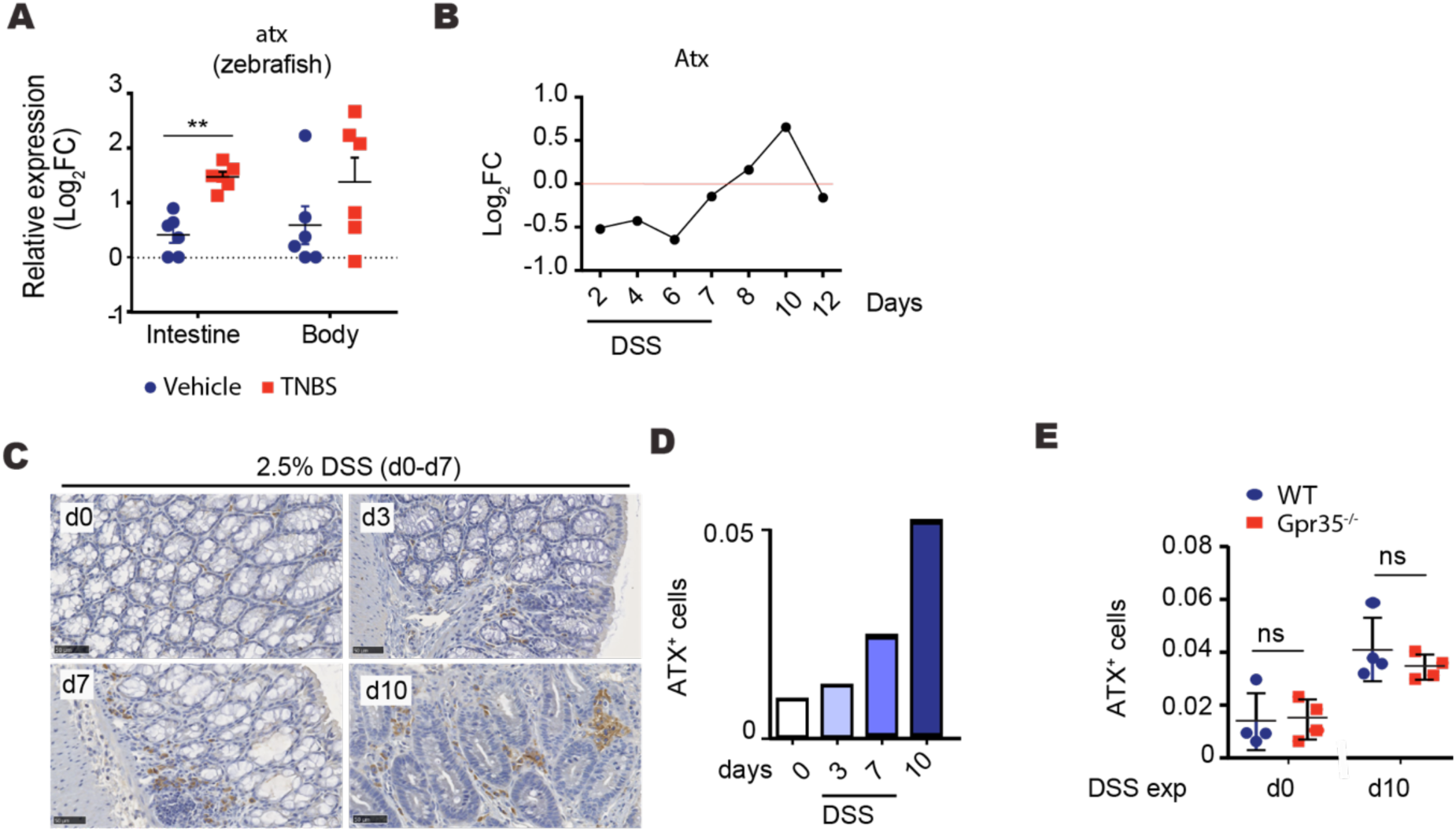
Colitis Induces Expression of the LPA-Generating Enzyme Autotaxin. (A) *Autotaxin (atx)* mRNA expression levels measured by qRT-PCR normalized to *ef1a* across the body and dissected intestine of WT zebrafish treated with TNBS or vehicle. Larvae were treated for 48 h starting at 72 hpf. Analysis was performed at 120 hpf and each dot represents a pool of 10 larvae. (B) RNA-seq analysis showing *Atx* gene expression from colonic tissue during 2.5% DSS-induced colitis (7 days exposure) and recovery. Dots show the average from three different mice per data point. (C) Immunohistochemistry imaging of WT mice treated with 2.5% DSS at indicated timepoints. Sections were stained for autotaxin (brown) and H&E (blue). One representative experiment is shown from two experiments. Scale bars, 50 μm. (D) Quantification of autotaxin^+^ cell data as shown in (C) from colonic tissue of DSS-treated mice at indicated time points. One representative experiment is shown from two experiments. (E) Quantification of autotaxin^+^ cells from colonic tissue at day 0 and day 10 of WT and *Gpr35*^-/-^ mice treated with DSS. Data are represented as individual values with mean ± SD. *p ≤ 0.05, **p ≤ 0.01, ***p ≤ 0.001 by two-way ANOVA with Tukey’s multiple comparisons test.

### Macrophage-Specific Deletion of *Gpr35* Exacerbates DSS Colitis

Given that GPR35 activation modulate cytokine production and macrophage migration together with the potential LPA synthesis during DSS-induced colitis, we hypothesize that GPR35 might affect intestinal inflammation. We next exposed WT and G*pr35*^-/-^ mice to DSS and evaluate the degree of intestinal inflammation. *Gpr35*-deficient mice had exacerbated colitis compared to WT animals as indicated by elevated body weight loss, increased disease activity scores, shorter colons, worsened histological signs of colitis, and more severe histology scores (Figures S5A-S5F). We next sought to define the cell type(s) underlying the worsened colitis in *Gpr35*-deficient mice. Since GPR35 is expressed by both intestinal epithelial cells and CX3CR1^+^ macrophages, we generated *Gpr35^flox^* mice by adding loxP sites before exon 2 and after 3′ UTR regions of *Gpr35* (Figure S6A) and crossed *Gpr35^flox^* with *Cx3cr1^CreER^* mice to obtain tamoxifen-inducible *Gpr35^ΔCx3cr1^* mice. This cross yielded mice with tamoxifen-inducible deletion of *Gpr35* specifically in CX3CR1^+^ macrophages, a deletion we confirmed by immunofluorescent staining for GPR35 (Figures S6B and S6C). *Gpr35^ΔCx3cr1^* mice displayed aggravated colitis compared to other control groups, as demonstrated by significant body weight loss, increased disease activity scores, significantly reduced colon length, and increased endoscopic and histological colitis scores (Figures 5A-5G). Consistent with enhanced inflammation, flow cytometric analysis revealed increased percentages and numbers of neutrophils in *Gpr35^ΔCx3cr1^* compared to control mice, further indicating exacerbated inflammation in the colon of *Gpr35^ΔCx3cr1^* mice compared to WT mice (Figures S7A-S7C). Aggravated colitis did not accompany higher frequencies or increased numbers of macrophages in the colonic lamina propria of *Gpr35^ΔCx3cr1^* mice (Figures S7A-S7C). *Gpr35^ΔCx3cr1^* mice did show reduced frequencies and mean fluorescence intensity of TNF-producing macrophages (Figures 6A and 6B), although we did not observe significant changes in overall expression of *Il10*, *Il1b*, *Il6*, or *Tnf* in colonic tissue (Figure S7D). These results indicate that deletion of *Gpr35* in macrophages exacerbates DSS-induced colitis and is associated with reduced macrophage-derived TNF.

**Figure 5.**
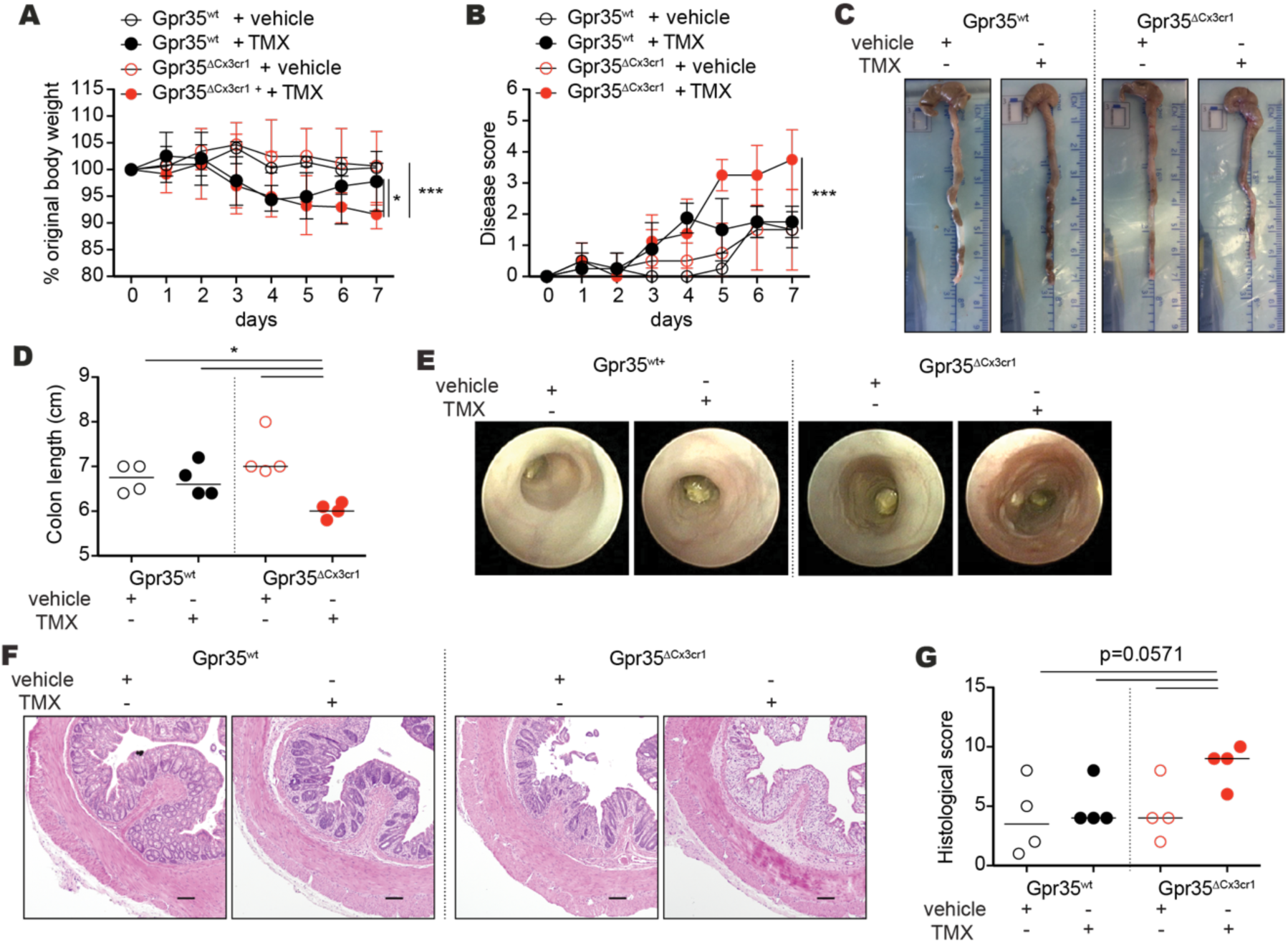
Deletion of *Gpr35* in CX3CR1^+^ Macrophages Exacerbates DSS-Induced Colitis. (A) Body weight changes (normalized to initial weight) during DSS colitis for 7 days from vehicle-injected (corn oil) or tamoxifen (TMX)-injected *Gpr35^wt^* or *Gpr35^ΔCx3cr1^* mice. Data are shown as mean ± SD for four mice per group. (B) Disease activity scores from daily monitoring of vehicle or TMX-injected *Gpr35^wt^* or *Gpr35^ΔCx3cr1^* mice with DSS colitis. Data are shown as mean ± SD for four mice per group. (C) Representative images of colons from vehicle or TMX-injected *Gpr35^wt^* or *Gpr35^ΔCx3cr1^* mice on day 7 of DSS colitis. (D) Colon lengths on day 7 of DSS colitis from DSS-treated *Gpr35^wt^* or *Gpr35^ΔCx3cr1^* mice treated with daily injections of vehicle or TMX. (E) Endoscopic images and (F) Representative H&E microscopy images of colons from vehicle or TMX-treated *Gpr35^wt^* or *Gpr35^ΔCx3cr1^* mice with DSS colitis. Scale bar, 100 μm. (G) Histology scores of indicated groups quantified from H&E staining of colon sections as shown in F. Each dot represents one animal with medians unless stated otherwise. *p ≤ 0.05, **p ≤ 0.01, ***p ≤ 0.001, ****p ≤ 0.0001 by two-way ANOVA with Tukey’s multiple comparisons test (A, B) or Mann-Whitney (D, G).

**Figure 6.**
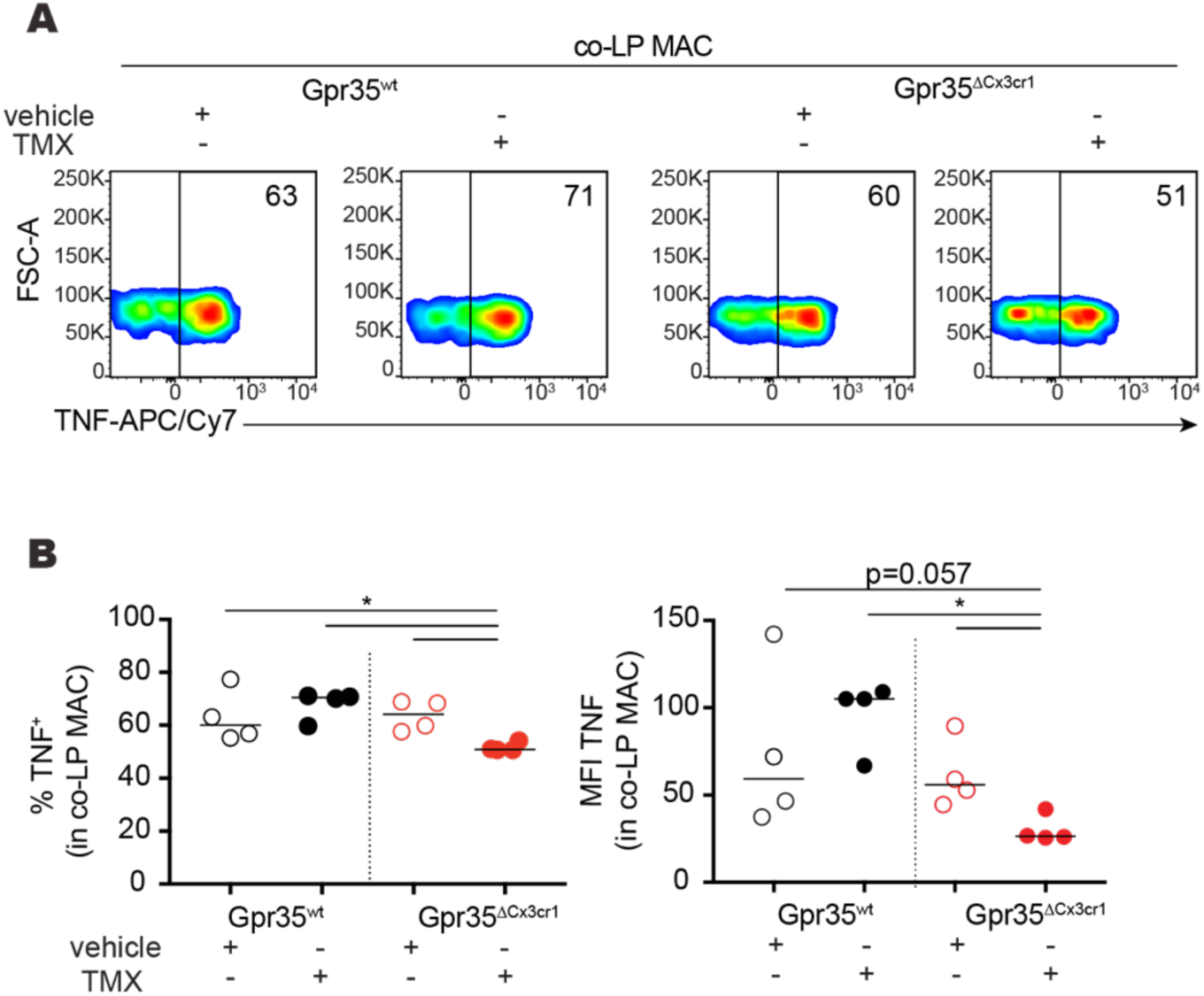
Deletion of *Gpr35* in CX3CR1^+^ Macrophages Is Associated with Reduced TNF Production. (A) After gating on viable CD45^+^MHC class II^+^ CD64^+^ cells from vehicle or TMX-treated *Gpr35^wt^* or *Gpr35^ΔCx3cr1^* mice TNF-producing colonic lamina propria macrophages (co-LP MAC) were analyzed by flow cytometry on day 7 of DSS colitis. Numbers in density plots indicate the percentage of TNF^+^ cells. (B) Percentages and mean fluorescence intensity (MFI) of TNF^+^ cells in macrophages from vehicle or TMX-treated *Gpr35^wt^* or *Gpr35^ΔCx3cr1^* mice based on flow cytometry data on day 7 of DSS colitis as shown in (A). Data are presented as individual values with medians; each dot represents one animal. *p ≤ 0.05, Mann-Whitney U test.

### TNF Attenuates Exacerbated DSS Colitis in *Gpr35*^ΔCx3cr1^ Mice

Despite the well-known pro-inflammatory properties of TNF and its pathogenic role in human IBD, some studies have suggested anti-inflammatory properties for TNF in the context of DSS-induced colitis, where TNF neutralization or *Tnf* deficiency exacerbates symptoms (Naito et al., 2003; Noti et al., 2010). We therefore investigated whether exacerbated colitis in *Gpr35^ΔCx3cr1^* mice was due to the inability of CX3CR1^+^ macrophages to produce TNF. To address this hypothesis, we injected *Gpr35^ΔCx3cr1^* mice daily with 1 μg TNF. Remarkably, we found that this treatment resulted in reduced colitis severity compared to untreated *Gpr35^ΔCx3cr1^* mice, as indicated by significantly decreased body weight loss, reduced disease activity scores, significantly longer colon length, reduced endoscopic signs of colitis, and reduced histologic colitis scores (Figures 7A-7F). Given that TNF can induce the expression of *Cyp11a1* and *Cyp11b1*, which encode steroidogenic enzymes involved in the synthesis of corticosterone in intestinal epithelial cells and thereby can attenuate the severity of DSS-induced colitis (Noti et al., 2010), we measured *Cyp11a1* and *Cyp11b1* expression in the colon in these mice. Expression of *Cyp11b1* but not *Cyp11a1* was reduced in *Gpr35^ΔCx3cr1^* mice (Figures 7G and 7H), and injection of TNF in *Gpr35^ΔCx3cr1^* mice restored *Cyp11b1* expression during colitis (Figure 7H). Consistent with these findings, supernatants of colonic explants from *Gpr35^ΔCx3cr1^* mice had lower corticosterone concentrations compared to WT animals with colitis, whereas TNF injection resulted in significantly increased corticosterone concentrations in supernatants of colonic explants compared to explants from untreated *Gpr35^ΔCx3cr1^* mice with colitis (Figure 7I). Taken together, these results demonstrate that loss of *Gpr35* in macrophages leads to aggravated colitis that is associated with reduced *Cyp11b1* expression in the intestine; notably, both the worsened colitis and decreased *Cyp11b1* expression were reversed by injection of TNF.

**Figure 7.**
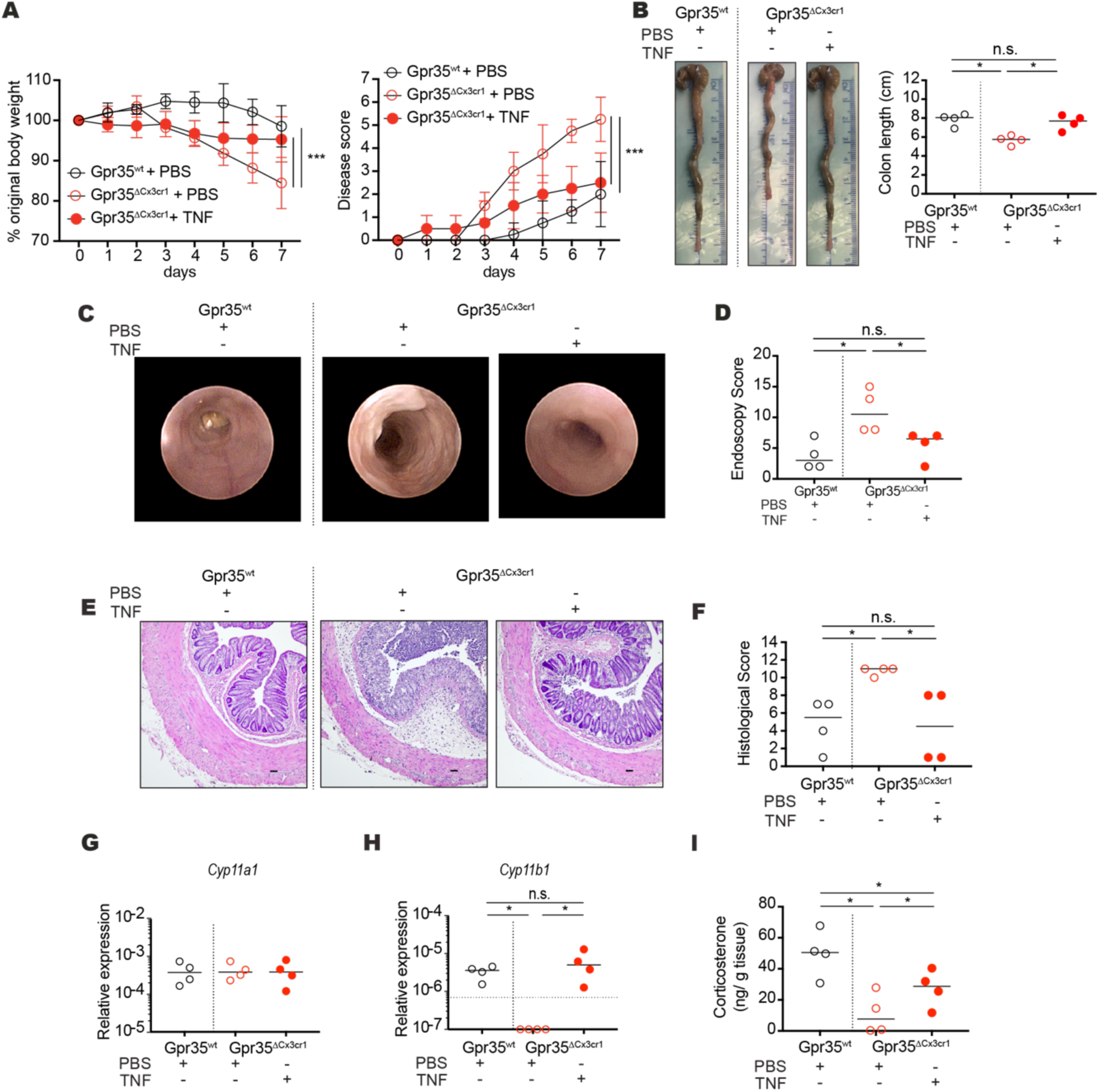
TNF Treatment of *Gpr35^ΔCx3cr1^* Mice Attenuates Colitis. (A) Body weight changes (normalized to initial weight) during 8 days of DSS exposure from PBS-injected *Gpr35^wt^* (WT) and PBS- or TNF-injected tamoxifen-treated *Gpr35^ΔCx3cr1^* mice. Data are shown as mean ± SD for four mice per group. Disease activity scores assessed by daily monitoring of DSS colitis of PBS-injected *Gpr35^wt^* and PBS- or TNF-injected tamoxifen-treated *Gpr35^ΔCx3cr1^* mice. Data are shown as mean ± SD for four mice per group. (B) Colon lengths on day 7 of DSS colitis from PBS-injected *Gpr35^wt^* (WT) and PBS- or TNF-injected tamoxifen-treated *Gpr35^ΔCx3cr1^* mice. (C) Endoscopic images and (D) Quantified endoscopic scores. (E) Representative H&E microscopy images of colon of PBS-injected *Gpr35^wt^* and PBS- or TNF-injected tamoxifen-treated *Gpr35^ΔCx3cr1^* mice on day 7 of DSS colitis. Scale bar, 50 μm. (F) histology scores. (G-H) mRNA expression levels of *Cyp11a1* (G) and *Cyp11b1* (H) relative to *Actb* by qRT-PCR in colon from PBS-injected *Gpr35^wt^* and PBS- or TNF-injected tamoxifen-treated *Gpr35^ΔCx3cr1^* mice on day 7 of DSS colitis. (I) Corticosterone concentration in supernatants of colonic explants of PBS-injected *Gpr35^wt^* and PBS- or TNF-injected tamoxifen-treated *Gpr35^ΔCx3cr1^* mice on day 7 of DSS colitisnormalized to weights of colonic tissues. Data are presented as individual values with medians. Each dot represents one biological replicate. *p ≤ 0.05, **p ≤ 0.01, ***p ≤ 0.001, ****p ≤ 0.0001 by two-way ANOVA with Tukey’s multiple comparisons test (A, B) or Mann-Whitney (, D, F, G, H, I).

## Discussion

It has been proposed that the recognition of host- and/or microbial-derived metabolites by GPCRs plays a critical role in driving cytokine responses that facilitate tissue destruction and resolution of inflammation in the context of infection or IBD (Chen et al., 2019; Cohen et al., 2017). In this study, we used a combination of genetic mouse and zebrafish models to study the biological relevance of GPR35 signaling, which dysfunction has been associated with increased IBD susceptibility. We have shown that LPA activates GPR35 leading to the regulation of the cytokine intestinal milieu at steady state condition as well as during intestinal inflammation. Furthermore, GPR35 expression distinguishes two distinct macrophage populations and is required in CX3CR1^+^ macrophages to trigger immunosuppressive pathways during intestinal inflammation.

Previous studies have suggested several potential ligands of GPR35, however depending on the experimental settings and/or species the results have been rather inconsistent (Binti Mohd Amir et al., 2018; Mackenzie et al., 2011; Maravillas-Montero et al., 2015; Oka et al., 2010; Southern et al., 2013). Because interspecies variation in ligand pharmacology must be considered for GPR35 (Milligan, 2018) we used genetic zebrafish and mouse models that lack GPR35. Using this comparative approach, we show remarkably conserved inflammatory cytokine production upon LPA stimulation, which was GPR35-dependent. Moreover, mouse and zebrafish macrophages responded to LPA as a chemoattractant in a GPR35-dependent manner. Although we demonstrated that GPR35 was required in bone marrow-derived macrophages in vitro, our in vivo zebrafish experiments did not allow us to rule out that other GPR35^+^ expressing cells sense LPA to then induce macrophage recruitment. Whether GPR35-dependent chemotaxis contribute to intestinal homeostasis and disease remains to be investigated. However, our findings suggest that LPA can activate GPR35 in zebrafish and mice, demonstrating that LPA-induced GPR35 signaling is conserved across species. We found that both mice and zebrafish showed increased expression of *Atx,* which catalyzes the formation of LPA (Gesta et al., 2002), during inflammation. However, the degree to which host cells and/or microbiota contribute to LPA production has not been fully explored. One possibility is that the disruption of epithelial cells leads to the release of LPA or precursors that are further metabolized by lamina propria cells expressing ATX. Phospholipids derived from microorganisms that constitute the intestinal microbiota may be another source of LPA during colitis (Cullinane et al., 2005). One recent study used mass spectrometry to distinguish microbial-derived versus host-derived metabolites by stable isotope tracing of ^13^C-labeled live non-replicating *E. coli* from ^12^C host isotypes (Uchimura et al., 2018). Our analysis of this published mass spectrometry data indicated that both host cells and microbes contribute to the LPA content in the colon. However, if LPA plays a role and the potential source during intestinal inflammation in IBD patients carrying the *GPR35* variant remains to be explored.

Antibiotic treatment of zebrafish and mice, in addition to expression analysis in germ-free mice, yielded insights into the modulation of GPR35 expression by the microbiota. Our results indicate that the microbiota modulates *Gpr35* expression in zebrafish and mouse. This microbiota dependence appeared to be specific to macrophages and was not observed in colonic epithelial cells, suggesting that the mechanism of *Gpr35* induction differs between cell compartments. Furthermore, the exposure of zebrafish to *V. anguillarum* indicated that pathogens may drive *gpr35* expression in zebrafish. This phenomenon was further confirmed in mice, in which *E. coli* induced the expression of *Gpr35* by CX3CR1^+^ macrophages, which are known to extend processes between epithelial cells to sample commensals, pathogens, and fungi (Leonardi et al., 2018; Niess et al., 2005; Rossini et al., 2014). Chemically induced intestinal inflammation also resulted in enhanced *Gpr35* expression as observed in zebrafish (TNBS-induced inflammation) and in mice (DSS-induced colitis). In agreement with the significant inflammation-induced increase of *Gpr35* expression in macrophages, *Gpr35*-deficient mice displayed more severe colitis compared to WT mice, and had reduced TNF production by macrophages.

TNF is a well-described pro-inflammatory cytokine that is central to the pathogenesis of Crohn’s disease and ulcerative colitis, as increased numbers of TNF-producing mononuclear cells are present in the lamina propria of patients with Crohn’s disease or ulcerative colitis (Reinecker et al., 1993). As a consequence, the inhibition of TNF with antibodies ameliorates colitis in animal models, and anti-TNF antibodies are essential for the treatment of patients with IBD (Corazza et al., 1999; Neurath et al., 1997; Present et al., 1999; Sands et al., 2001; Siegel et al., 1995). Our data suggest a model in which GPR35 signaling in macrophages induce TNF expression which is beneficial to maintain intestinal homeostasis, which might be in contradiction with the paradigm that TNF is pathogenic in IBD. In this line, some studies have shown that TNF has anti-inflammatory effects in the context of *Tnf*-deficient animals which result in more severe DSS-induced colitis (Naito et al., 2003; Noti et al., 2010). TNF is quickly released after tissue damage to reduce damage-associated mortality (Mizoguchi et al., 2008). Our data show that GPR35-dependent TNF induction result in induction of corticosterone production, which might in turn suppress immune responses. Therefore, TNF can play a protective or deleterious role depending on the context and stage of the disease. On the other hand, studies in macrophages carrying the IBD-associated T108M polymorphism in GPR35 result in enhanced metabolic activity compared to the wild type GPR35, which is the opposite effect compared to GPR35 loss of function (Schneditz et al., 2019; Tsukahara et al., 2017). This data suggests the possibility that IBD patients carrying the T108M polymorphism might have enhanced TNF production, which result in aberrant inflammatory immune responses.

We cannot exclude the possibility that other endogenous ligands may bind to GPR35 and that LPA may also modulate the intestinal cytokine milieu by binding to other LPA receptors. Before GPR35 can be considered as a possible target in the clinic for the treatment of IBD, better pharmacological screenings must be considered to identify additional putative ligands and inter-species variations of potential ligands. The identification of possible connections between host- and microbial-derived metabolites with the immune system will be critical in the future to dissect mucosal immune responses in healthy individuals and during colitis.

## Acknowledgments

This work is part of the Ph.D. thesis of B.K. Philippe Demougin, Biozentrum, Basel. Christian Beisel, Genomics Facility Basel, ETH Zürich, helped with RNA-seq. Florian Geier, Bioinformatics Core, Department of Biomedicine, University of Basel, helped with the analysis of the RNA-seq data. Calculations were performed at the sciCORE (http://scicore.unibas.ch/) scientific computing center at the University of Basel. We thank all gastroenterologists who enrolled patients into the Swiss IBD Cohort. J.H.N. was supported by SNSF grants 310030_175548 and 316030_170809. T.K. was supported by SNSF M.D. Ph.D. fellowship 323530_183981; the Swiss IBD Cohort Investigators were supported by SNSF grant 33CS30-148422. E.J.V. was supported by grants from the Swedish Research Council, VR grant K2015-68X-22765-01-6, Formas grant nr. FR-2016/0005, and the Wallenberg Academy Fellow program.

## Author Contributions

B.K., C.D.C., Ph.W., O.E.D., R.A.M., H.M., T.K., S.D., and C.K.A. performed experiments. P.H. provided patient samples, and P.P.H provided *V. anguillarum* extracts. E.J.V. and J.H.N. conceived the idea. B.K., C.D.C., E.J.V., and J.H.N wrote the paper. All authors discussed the data, read, and approved the manuscript.

## Material and Methods

### Mouse lines

C57BL/6, *Rag2*^-/-^, *Cx3cr1*-GFP (B6.129P-Cx3cr1^tm1Litt/J^) and Cx3cr1^CreER^ (B6.129P2(Cg)-*Cx3cr1^tm2.1(cre/ERT2)Litt^*^/W gan J^) mice were bred and maintained in the animal facility of Department of Biomedicine, University of Basel, Switzerland or the respective facility at the Karolinska Institutet, Solna, Sweden. *Gpr35*-tdTomato, *Gpr35*^-/-^ and *Gpr35^flox/flox^* animals were constructed as described below. *Gpr35*-tdTomato mice were crossed with *Cx3cr1*-GFP mice to generate double reporter mice, and *Gpr35^flox/flox^* were crossed with *Cx3cr1^CreER^* to obtain *Gpr35^ΔCX3CR1^* mice, in which the tamoxifen-inducible, Cre-mediated recombination will lead to the excision of GPR35 in CX3CR1^+^ cells. All animals were kept under specific pathogen-free (SPF) conditions. Germ-free C57BL/6 mice were obtained from the Core Facility for Germ-Free Research at the Karolinska Institutet, Solna, Sweden. For in vivo and in vitro experiments at least 3 mice per group were included. Animals between 6-12 weeks of age were randomly selected for experimental groups. All mouse experiments were conducted in accordance to the Swiss Federal and Cantonal regulations (animal protocol number 2832 (canton Basel-Stadt)) and the Stockholm regional ethics committee under approved ethical number N89-15.

### Generation of *Gpr35*-IRES-tdTomato knock-in mice

*Gpr35*-IRES-tdTomato knock-in mouse line was generated by Beijing Biocytogen (Beijing, China) by introducing IRES-tdTomato between the protein coding sequences of the targeted gene and 3’UTR under the genetic background of C57BL/6J. In brief, for construction of the targeting vector, 4.7-kb left homology arm spanning exon 1 and an FRT-flanked neo cassette were inserted 352bp upstream of exon 2; an internal ribosome entry site 2 (IRES2) sequence (allows translation initiation in the middle of an mRNA sequence), a tdTomato reporter and 3.9-kb right homology arm were inserted just downstream of stop codon. The complete sequence of the targeting vector was verified by sequencing analysis. After linearization, the targeting vector was transfected into C57BL/6J embryonic stem (ES) cells by electroporation. Eight positive ES clones were identified by Southern blot analysis with 5’probe and 3’probe, and Karyotype analysis. Positive ES clones were injected into BALB/c blastocysts and implanted into pseudopregnant females. Four chimeric male mice were crossed with FLP females to obtain F1 mice carrying the recombined allele with the removal of Neo selection cassette. The F1 mice were validated for germinal line transmission of the recombination event by using the PCR strategy. The elimination of the neo cassette in the offspring was analyzed by PCR with the primers Frt-F2 and Frt-R2 (Table S1). Male and female heterozygous mice were crossed to produce homozygous mutant mice. Reporter animals were genotyped by PCR with primers listed in Table S3. Following PCR cycling parameters were used with 35 cycles of amplification: denaturation 95°C for 2 min; amplification 95°C 30 sec, 62°C 30 sec, 72°C 25 sec; final elongation 72°C 10 min.

### Generation of Gpr35-flox and knock-out (KO) mice

*Gpr35^flox^* and *Gpr35^-/-^* mice were generated using the CRISPR/Cas9 system by Beijing Biocytogen (Beijing, China). Briefly, the Cas9/guide RNA (gRNA) target sequences were designed to the regions upstream of exon2 and downstream of 3’UTR. The targeting construct of *Gpr35^flox^* consisting of 1.3 kb arms of homologous genomic sequence immediately upstream (5’) of exon 2 and downstream (3’) of 3’UTR flanked by two loxP sites (Figure S3A). Cas9 mRNA and sgRNAs were transcribed with T7 RNA polymerase in vitro. Cas9 mRNA, sgRNAs and donor vector were mixed at different concentrations and co-injected into the cytoplasm of fertilized eggs at the one-cell stage. The genotypes for *Gpr35^flox^* and *Gpr35^-/-^* mice were validated by PCR amplification and direct sequencing. *Gpr35^flox^* mice were further validated by Southern blot analysis.

For *Gpr35* targeting, two sgRNAs were designed to target the regions upstream of exon 2 and downstream of 3’UTR. For each targeted site, candidate sgRNAs were designed using the CRISPR design tool (http://www.sanger.ac.uk/htgt/wge/). sgRNAs were screened for on-target activity using the UCA kit (Lin et al., 2016). Cas9 mRNA and sgRNAs were transcribed with T7 RNA polymerase in vitro. For Cas9 mRNA and sgRNA production, the T7 promoter sequence was added to the Cas9 and sgRNA templates by PCR amplification. T7-Cas9 and T7-sgRNA PCR products were gel purified and used as the template for in vitro transcription with the MEGAshortscript T7 kit (Life Technologies) according to the kit protocol. Cas9 mRNA and sgRNAs were purified using the MEGAclear kit and eluted with RNase-free water. The targeting construct of Gpr35 flox consisting of 1.3 kb arms of homologous genomic sequence immediately upstream (5’) of exon 2 and downstream (3’) of 3’UTR flanked by two loxP sites (Figure S3C). The donor vector was prepared using an endotoxin-free plasmid DNA kit. C57BL/6N females were used as embryo donors and pseudopregnant foster mothers. Superovulated C57BL/6N mice (3-4 weeks old) were mated to C57BL/6N stud males, and fertilized embryos were collected from the ampullae. Cas9 mRNA, sgRNAs and donor vector were mixed at different concentrations and co-injected into the cytoplasm of fertilized eggs at the one-cell stage. After injection, surviving zygotes were transferred into the oviducts of KM pseudopregnant females. The genotyping of *Gpr35*-deficient animals was done by PCR in 2 different reactions using the listed primers (Table S3) under the following conditions: initial denaturation at 95°C for 3 min; 35 cycles of denaturation 95°C 30 sec, annealing 64°C 30 sec, elongation 72°C 45 sec; and final elongation 72°C 10 min. The *Gpr35*-flox mice were genotyped by PCR (for primers see Table S3) by denaturating at 95°C for 3 min, amplifying 35 cycles at 95°C 30 sec, 62°C 30 sec, 72°C 35 sec and elongating at 72°C for 10 min.

### Zebrafish lines

The *Tg(mpeg1:mCherry)* was kindly provided by Professor Georges Luftalla (Montpellier, France). The zebrafish predicted gene G-protein coupled receptor 35-like (LOC101882856) (mRNA sequence ID: XM_021466387.1, previous Ensembl ID: ENSDARG00000075877, current Ensembl ID: ENSDARG00000113303) was targeted using a CRISPR-Cas9 approach by the Genome Engineering Zebrafish, Science for Life Laboratory (SciLifeLab), Uppsala, Sweden. CRISPR/Cas9 gene editing was performed as previously described (Li et al., 2016) and the gRNA was targeted within exon 2 in the reverse strand with a gene specific gRNA-target sequence followed by a protospacer adjacent motif (PAM), (5’ GGT AGG CCA CAC GCT CAA AC<u>A GG</u> 3’ – PAM sequence is underlined). Eggs from WT AB strain were co-injected with a total volume of 2nL consisting of a mix of 300 pg Cas9 mRNA and 25pg of sgRNA at the single-cell stage. Founder screening by somatic activity test (CRISPR-STAT) and germline transmission were assayed using fluorescence PCR as previously described (Li et al., 2016). Briefly, injection groups with somatic activity were grown to adulthood for founder screening and positively identified founders (F_0_) were in-crossed with another founder to screen for germline transmission in F_1_ embryos. F_1_ embryos were raised to adulthood, fin clipped and genotyped using fluorescence PCR followed by subsequent validation of the mutation using Sanger sequencing. F_1_ heterozygotes were outcrossed with AB fish and the resulting F_2_ heterozygotes were further maintained and in-crossed. The F_3_ embryos were raised to adulthood and screened for homozygous mutants and wild type zebrafish by PCR based genotyping (WT forward primer: 5’-TAG CCT GTT TGA GCG TGT GG-3’; mutant forward primer: 5’-CCA TTA GCC TGT GGC CT -3’; common reverse primer: 5’-CAC CAG CGA TTT GGT CAG AA-3’), which were further in-crossed (i.e. ‘mutant with mutant’ and ‘wildtype with wildtype’) to generate mutant and WT embryos that were subsequently used for experiments. For the purpose of experiments, the mating was performed in a random fashion at all occasions. For husbandry, embryos were kept and raised to adulthood in systems with circulating, filtered and temperature (28.5 °C) controlled water. All procedures were performed according to Swedish and European regulations and have been approved by the Uppsala University Ethical Committee for Animal Research (C161.14) and Karolinska Institutet Ethical Committee for Animal Research (N5756/17). <u>Primers used for fluorescence PCR:</u> Forward M13F-tailed primer: 5’-TGT AAA ACG ACG GCC AGT CTC AAG CAA ACT GCT TCC TCT T-3’; Reverse PIG-tailed primer: 5’-GTG TCT TGC ATG TAG ATG TGA GTG TCG GT-3’; M13F FAM primer: /56FAM/ TGT AAA ACG ACG GCC AGT

### Human inflammatory bowel disease biopsies

The study population for mRNA analysis included 31 patients with Crohn’s disease and 31 patients with ulcerative colitis (20 with active disease, 11 in remission) recruited to Swiss Inflammatory Bowel Disease Cohort Study (SwissIBD cohort project 2016-12) started in 2006 (Pittet et al., 2009). The diagnoses of Crohn’s disease and ulcerative colitis were validated by endoscopy, radiology or surgery at least 4 months before recruitment of the patients. Patients with colitis or ileitis caused by other conditions or with no permanent residency in Switzerland were excluded from the study. Ileocolonoscopy was done to confirm quiescent IBD or to determine the activity in active IBD. For active IBD, biopsies were taken from macroscopically inflamed regions. Table S2 gives detailed depiction of patient information. After the collection, the biopsies were kept in RNAlater® stabilization solution (Invitrogen) at -80°C until further use. The study population for immunofluorescence involved 4 ulcerative colitis and 3 Crohn’s disease patients recruited to the Basel IBD cohort. The biopsies were taken from inflamed or non-inflamed regions of the same patients following ileocolonoscopy. The specimens were embedded in optimal cutting temperature (OCT) compound (TissueTek) and stored at -80°C. Table S3 shows the detailed patient characteristics (ethics protocol EKBB 139/13 (PB 2016.02242) (Ethics Committee for Northwest and Central Switzerland (EKNZ).

### Cell lines

The human colon adenocarcinoma HT29 (ATCC, HTB-38) cell line was cultured in Dulbecco’s Modified Eagle Medium (DMEM)-GlutaMAX (Invtirogen) with 10% FBS (Invitrogen), 100 U/mL penicillin and 0.1 mg/mL streptomycin (Invitrogen). The cells were incubated at 37°C, 5% CO_2_ and medium were changed every 3 days and cells were passaged with 0,05% trypsin-EDTA (Invitrogen) twice per week.

### Dextran sodium sulfate induced colitis mouse model

Weight-matched 6 to 12-week-old female mice were administered with 1.5-2.5% DSS (MP Biomedicals) in the drinking water for 5 days followed by 2 days of normal drinking water. Mice were daily weighed and monitored for clinical colitis score. Clinical colitis scores were calculated according to the following criteria (Steinert et al., 2017): rectal bleeding: 0 - absent, 1 - bleeding; stool consistency: 0 - normal, 1 - loose stools, 2 - diarrhea; position: 0 - normal movement, 1 - reluctant to move, 2 - hunched back; fur: 0 - normal, 1 - ruffled, 2 - spiky; weight loss: 0 – no loss, 1 - body weight loss 0-5%, 2 - body weight loss >5 - 10%, 3 - body weight loss > 10 - 15%, 4 - body weight loss > 15%. Endpoints of the experiment are total score of ≥ 6, > 15 % body weight loss, excessive bleeding, and rectal prolapse.

### Hematoxylin-eosin (H&E) staining and histological scoring

5 µm paraffin sections from mouse colon were stained with H&E. Histological scores for colonic inflammation were assessed semi-quantitatively using the following criteria (Souza et al., 2017; Steinert et al., 2017): mucosal architecture (0:normal, 1-3: mild-extensive damage); cellular infiltration (0:normal, 1-3: mild-transmural); goblet cell depletion (0:no, 1:yes); crypt abscesses (0:no, 1:yes); extend of muscle thickening (0: normal, 1-3: mild-extensive). Tissues were scored by at least two blinded investigators and data is presented by the mean.

### Mouse Endoscopy

To assess macroscopic colitis severity, mice were anaesthetized with 100 mg/kg body weight ketamine and 8 mg/kg body weight Xylazine intraperitoneally. The distal 3 cm of the colon and the rectum were examined with a Karl Storz Tele Pack Pal 20043020 (Karl Storz Endoskope, Tuttlingen, Germany) as previously described (Melhem et al., 2017).

### Treatment of zebrafish with 2,4,6-Trinitrobenzenesulfonic acid or with antibiotics

To induce inflammation, zebrafish larvae were either untreated or treated with 2,4,6- Trinitrobenzenesulfonic acid (TNBS; Sigma Aldrich P2297) from day 3 post-fertilization until 120 hpf. TNBS was added in a 1:1000 dilution in E3 water (final concentration: 50 µg/mL) and replaced every 24 hours. To deplete the bacterial content, zebrafish larvae were treated with an antibiotic cocktail from day 3 post-fertilization until 120 hpf (Bates et al., 2006). The antibiotic cocktail consists of Ampicillin (100 µg/ml) and Kanamycin (5 µg/ml) that was added to E3 water and replaced every 24 hours.

### Treatment of mice with antibiotics

WT mice were treated with antibiotic cocktail for 10 consecutive days by oral gavage. The antibiotic cocktail contains Ampicillin (1mg/ml), Kanamycin (1mg/ml), Gentamicin (1mg/ml), Metronidazole (1mg/ml), Neomycin (1mg/ml), and Vancomycin (0.5mg/ml).

### Preparation of sense and antisense Digoxigenin (DIG)-labeled RNA probes for detection of *Gpr35b* in zebrafish

DNA plasmid containing *Gpr35b* cDNA (5 µg) was linearized in a 2 h digestion, using SacI and EcoRI to generate the sense and anti-sense probe template, respectively. The linearized plasmid was purified by phenol: chloroform extraction method followed by ethanol precipitation. Following successful production of template, the in vitro synthesis of the sense and antisense DIG-Labeled RNA probes were made in a 2 hours incubation at 37 °C with the following transcription mix (20 µL): DNA template (1-2 µg), DIG-RNA labeling mix, protector RNase Inhibitor, transcription buffer and RNA Polymerase T7 and T3, respectively. Following the incubation, DNA template was digested by adding DNase I for 30 min at 37 °C and was stopped by adding 2 μl of 0.2 M EDTA. DIG-Labeled RNA Probes were precipitated by LiCl method and resuspended in 30 μl Probe solution (19 μl sterilized water, 10 μl RNAlater and 1 μl 0.5M EDTA).

### In situ hybridization for *gpr35b* detection in zebrafish

*In situ* hybridization (ISH) was performed in whole zebrafish larvae from the developmental stages 72 hpf, 96 hpf and 120 hpf. Those larvae were fixed by 4% paraformaldehyde (PFA) in PBS at 4 °C overnight followed 3 PBS washes. Progressive dehydration by washing for 5 min in 25%, 50% and 75% methanol in PBS and final 5- and 15-min wash in 100% methanol were performed. In some cases, the depigmentation method was required due to their developmental stage. In this case, larvae were treated with 3% H_2_O_2_/0.5% KOH at RT until pigmentation has completely disappeared and then progressive dehydration was performed as described above. Larvae were placed at -20 °C for at least 2 h. After incubation, larvae were rehydrated, washed 4 times with PBST (0.1% Tween-20 in PBS) followed by proteinase K (10 µg/mL) treatment at RT for a time defined by the developmental state such is indicated coming up next: 72 hpf – 20 min; 96 hpf – 30 min; and 120 hpf – 40 min. Proteinase K digestion was stopped by incubating the larvae for 20 min in 4% PFA. Larvae were washed with PBST and prehybridized with 700 µL hybridization mix (HM) solution (50% deionized formamide (Millipore); 0.1% Tween-20 (Sigma); 5X saline sodium citrate solution (Merck); 50mg/ML of heparin (Sigma); 500 mg/mL RNase-free tRNA (Sigma)) for 5 h at 70 °C. HM solution was replaced by 200 µL of HM containing 50 ng of antisense/sense DIG-labeled RNA probe and incubated overnight at 70 °C. Then the larvae went through several washing steps with SSC and PBST solution followed by incubation with blocking buffer for 4h at RT. Afterward, larvae were incubated with anti-DIG-AP antibody solution overnight at 4 °C. Subsequently, they were washed 6 times for 15 min with gentle agitation on a horizontal shaker, incubated with alkaline Tris buffer for 5 min at RT with gentle agitation, and stained in dark using 700 μL staining solution. When the color was developed, the reaction was stopped by adding stop solution (1mM EDTA and 0.1% Tween-20 in PBS pH 5.5). Finally, larvae were transferred to a tube containing 100% glycerol and kept in this solution at least 24 h before mounting them.

### Challenging of mice with *E. coli*-CFP

*E. coli* DH10B pCFP-OVA was constructed as previously described (Rossini et al., 2014). Gpr35-tdTomato x Cx3cr1-GFP mice were gavaged every other day for 21 days with 1×10^8^ CFUs of CFP-OVA^+^ *E. coli* and sacrificed for further analysis.

### Exposure of zebrafish with *Vibrio anguillarum*

*V. anguillarum* strain 1669 was grown in tryptic soy broth medium to OD_600_ (optical density at 600 nm) 1.5. Bacterial pellet (9 ml of full-grown culture) was resuspended in NaCl (9 g/L), 0.35% formaldehyde, and incubated overnight at 20°C. The suspension was washed four times in NaCl(9 g/L) and resuspended in 800 ml of the same isotonic solution. *V. anguillarum* extract was mixed in a 1:1 ratio with phenol red (Sigma Aldrich P0290). One µL of this mixture was diluted with 2 µL PBS from which 2 nL were used to be injected in the intestinal lumen of 110 hpf zebrafish larvae. Larvae were anesthetized using 0.0016% Tricaine MS0222 (Sigma-Aldrich E10521). Larvae were then monitored for recovery and analyzed 6 h post injection.

### LPA injection in zebrafish

LPA (10 µM) or equal volume of DMSO were mixed with FITC-Dextran (500 µg/ml) in PBS. For the challenge, 2 nl were injected in the otic vesicle of 110 hpf larvae anesthetized with 0.0016% Tricaine MS-222. Larvae were then monitored for recovery and macrophage recruitment was analyzed 6 h after the injection.

### Stimulation of zebrafish larvae with LPA

WT and *gpr35b^uu1892^* zebrafish larvae were either left unstimulated or stimulated with 10 μM LPA (Sigma L7260) in water from 96 hpf until 120 hpf. After the incubation, zebrafish larvae were lysed, RNA extracted, and cytokine production was evaluated by qPCR using primers listed in Table S3.

### Cell Isolation from the small and large intestinal lamina propria, mesenteric lymph nodes and spleen

Colonic lamina propria cells were isolated as described previously (Radulovic et al., 2019) (Steinert et al., 2017). Briefly, extracted colon or small intestine segments were opened longitudinally and washed with PBS (Sigma-Aldrich). IECs were dissociated using 5 mM EDTA at 37 ^°^C in a shaking water bath at 200 rpm for 10 minutes. The dissociation step was repeated in fresh EDTA solutions for 2 additional times. The tissue was vortexed for 30 sec before and after each incubation and IECs were collected for further processing, if necessary. After removing IECs, the tissue was immersed in PBS to wash the EDTA away and cut into small pieces for digestion. The tissue was digested in Roswell Park Memorial Institute (RPMI) 1640 (Sigma-Aldrich) containing 0.5 mg/ml Collagenase type VIII (Sigma-Aldrich) and 10 U/mL DNase (Roche) for 15-20 min at 37 ^°^C in shaking water bath with 30 sec vortexing each 5 min. Digested tissue was passed through a 70 μm cell strainer and single cell suspension was pelleted for further analysis. Spleen and MLN cells were isolated by mashing the tissue with a syringe plunger on a 70 μm cell strainer. Spleen red blood cells (RBCs) were lysed using ammonium-chloride-potassium buffer (150 mM NH_4_Cl, 10 mM KHCO_3_, 0.1 mM 0.5 M EDTA). Remaining cells were pelleted for further use.

### Antibodies, cell staining and flow cytometry

Up to 5×10^6^ isolated cells were incubated for 30 min at 4 ^°^C with anti-CD16/CD32 (Fc receptor) clone 93 (Invitrogen) to block non-specific binding and with fixable viability dye eFluor455UV (eBioscience) for live/dead cell exclusion. Cells were washed in PBS containing 2% Fecal Bovine Serum (FBS), 0.1% sodium azide, and 10 mM EDTA (FACS buffer) and stained for surface antigens for 20 min at 4 ^°^C. For intracellular staining, cells were further fixed and permeabilized in Cytofix/Cytoperm solution according to the manufacturer’s instructions (BD Biosciences) followed by incubation with antibodies against intracellular antigens for 20 min at 4 ^°^C. Cells were then resuspended in FACS buffer and flow cytometric analysis was performed on a Fortessa flow cytometer (BD Biosciences). Data was analyzed using FlowJo software version 10.0.7r2 (TreeStar). In all experiments, doublet discrimination was done on forward scatter (FSC-H) versus FSC-A plot. Mononuclear phagocyte staining was done using antibodies eVolve655-conjugated anti-CD45 clone 30-F11 (eBioscience), Biotin-labeled anti-CD3 clone 145-2C11, anti-CD19 clone 6D5 and anti-NK.1.1 clone PK136, AF700-conjugated anti-I-A/I-E (MHCII) clone M5/114.15.2, PE/Cy7-conjugated anti-CD64 clone X54-5/7.1, APC/Cy7-conjugated anti-CD11c clone N418, FITC-conjugated anti-CD11b clone M1/70, PerCP/Cy5.5-conjugated anti-Ly6C clone HK1.4, and APC-conjugated anti-Ly6G clone 1A8 (all BioLegend). For lineage exclusion, CD3^+^, CD19^+^ and NK1.1^+^ cells were gated out. For lymphocyte staining, antibodies for APC/Cy7-conjugated anti-CD45 clone 30-F11, AF700-conjugated anti-CD3 clone 17A2, BV785-conjugated anti-CD19 clone 6D5, BV510-conjugated anti-CD4 clone RM4-5 and PerCP-conjugated anti-CD8a clone 53-6.7 or Biotin-labeled anti-CD8 clone 53-6.7 (all BioLegend) were used. For innate lymphoid cell panel, antibodies APC/Cy7-conjugated anti-CD90.2 clone 30-H12 (BioLegend), APC-conjugated anti-GATA3 clone 16E10A23 (BioLegend), PerCP/Cy5.5-conjugated anti-RORγT clone Q31-378 (BD Biosciences), PE/Cy7-conjugated anti-T-bet clone 4B10 (BioLegend) and FITC-conjugated anti-Eomes clone WD1928 (Invitrogen) were included whereas Biotin-conjugated antibodies anti-CD3 145-2C11, anti-CD8a 53-6.7, anti-CD19 6D5, anti-CD11c N418 (all BioLegend), anti-B220 RA3-6B2 (BD Biosciences), anti-Gr-1 RB6-8C5, anti-TCRβ H57-597, anti-TCRγδ GL3 and anti-Ter119 TER-119 (all BioLegend) were used for lineage exclusion. eFluor450 conjugated Streptavidin (eBioscience) was used for all biotin labeled antibodies.

### RNA extraction and quantitative PCR

RNA was extracted from cells, mouse or zebrafish tissues, whole zebrafish larvae or human biopsies using TRI Reagent (Zymo Research) or TRIzol (Invitrogen) according to the manufacturer’s instructions. For DSS-treated mouse colonic tissue, Direct-zol RNA MiniPrep kit (Zymo Research) was used to remove the DSS residues. RNA samples were treated with TURBO DNase (Invitrogen) and reverse transcribed using High Capacity cDNA Reverse Transcription (Applied Biosystems) or iScript cDNA synthesis (Bio-Rad) kits by following manufacturer’s instructions. Quantitative PCR was performed using primers listed in Table S3 and QuantiNova SYBR Green PCR (Qiagen) or iTaq™ Universal SYBR® Green Supermix (Bio-Rad) kits. Samples were run on an ABI ViiA 7 cycler or a CFX384 Touch Real-Time PCR. Ct values were normalized to that of *efa1*, *Hrpt*, *Gapdh* or *Actb* and relative expression was calculated by the formula 2^(-ΔCt). Used primers are listed in Table S1.

### Immunofluorescence staining

Human biopsies were provided by the Basel IBD cohort in cryoblocks. Mouse tissues were fixed with 4% PFA and left in 30% sucrose overnight for cryo-embedding or dehydrated in ethanol solutions for paraffin embedding. All tissues were sectioned at 6 µm and fixed with 4% PFA. Blocking and permeabilizing were done using PBS containing 0.4% Triton X-100 for cryosections or 0.1% Tween20 for paraffin sections and 5% goat serum (all Sigma-Aldrich). Tissue sections were stained with rabbit polyclonal anti-human/mouse GPR35 primary antibody and goat anti-rabbit IgG secondary antibody. For all samples, NucBlue^TM^ Live Cell Stain (Thermo Fisher) was used for nuclear staining and samples were imaged using a Nikon A1R confocal microscope.

### Autotaxin staining

Mouse tissues were fixed with 4% PFA and dehydrated in ethanol solutions for paraffin embedding. All tissues were sectioned between 5-6 µm. Endogenous peroxidase activity was blocked with 3% H_2_O_2_ solution in methanol and antigen retrieval to unmask the antigenic epitope was performed with EDTA buffer (1mM EDTA, pH 8.0). Blocking was done using BLOXALL™ Blocking Solution (Vector Laboratories SP-6000). Tissue sections were stained with mouse monoclonal anti-ENPP2 (autotaxin) Mouse/Human primary antibody (Abcam ab77104) and goat anti-rabbit IgG secondary antibody. All samples were additionally stained with H&E as described previously.

### *Ex vivo* imaging of colonic tissues

Extracted colon was washed with PBS, opened longitudinally and placed on a slide. A drop of PBS was added to prevent the tissue from drying and tissue was covered with a coverslip.

Tissues were imaged on a Nikon A1R confocal microscope.

### RNA sequencing

RNA was isolated from sorted GPR35^+^ and GPR35^-^ colonic macrophages from 1 or 2 *Gpr35*-tdTomato mice. RNA quality control was performed with an Agilent 2100 Bioanalyzer and the concentration was measured by using the Quanti-iT RiboGreen RNA assay Kit (Life Technologies). cDNA was prepared using SMART-Seq v4 Ultra Low Input RNA Kit (Tamara). Sequencing libraries were prepared using Nextera XT DNA Library Preparation Kit (Illumina). Indexed cDNA libraries were pooled in equal amounts and sequenced SR81 with an Illumina NextSeq 500 Sequencing system. Reads were aligned to the mouse genome (UCSC version mm10) with STAR (version 2.5.2a) using the multi-map settings ‘— outFilterMultimapNmax 10 --outSAMmultNmax 1’ (Dobin et al., 2013). Read and alignment quality was evaluated using the qQCReport function of the R/Bioconductor package QuasR (R version 3.4.2, Bioconductor version 3.6) (Gaidatzis et al., 2015). Assignment of reads to genes employed the UCSC refGene annotation (downloaded 2015-Dec-18). QuasR function qCount function was used to count the number of read (5’ends) overlapping with the exons of each gene assuming an exon union model. Differential gene expression analysis was performed using the R/Bioconductor package edgeR (McCarthy et al., 2012). After filtering genes with logCPM>1 in at least 1 sample, a paired design analysis was performed taking group and animal ID into account. Differential expression statistics for the GPR35_Plus vs. GPR35_Minus group contrast employed the glmQLFit and glmQLFTest functions of edgeR. Resulting P-Values were false discovery rate adjusted. RNA-seq data shown in Figure 4B has been obtain from a dataset published elsewhere (Czarnewski et al., 2019).

### 3’-5’-Cyclic adenosine monophosphate (cAMP) assay

To screen potential GPR35 ligands, the cAMP Hunter^TM^ eXpress assay platform (Eurofins) was used according to the manufacturer’s directions. Briefly, GPR35-transfected CHO-K1 cells were thawed and 3×10^5^ cells were seeded on a 96-well plate followed by overnight incubation at 37 ^°^C, 5% CO_2_. Cells were treated at 37 ^°^C for 30 minutes with 15 μm Forskolin and 1:3 serial dilutions of potential ligands with the following starting concentrations: 10 μM recombinant human CXCL17 (R&D Systems), 10 μM lysophosphatidic acid (LPA) or 10 mM kynurenic acid (KYNA) (both Sigma-Aldrich). Zaprinast (Sigma-Aldrich) was used as a positive control. cAMP levels were measured by enzyme-fragment complementation (EFC) technology where two fragments of β-galactosidase were used. In presence of cAMP, cAMP labelled with one part of the enzyme is outcompeted to bind to anti-cAMP antibody and therefore is free to complement the enzyme complex and cleave the substrate to produce a luminescent signal. The signal was then measured by Synergy H1 Microplate Reader (Biotek).

### Mouse bone marrow-derived macrophages

Femur and tibias from WT or *Gpr35*-deficient mice were cut at both ends and bone marrow was flushed out with PBS with the help of a syringe with 25-gauge needle. The cells were collected and cultured in RPMI 1640 containing 10% FBS, 0.05 mM 2-ME, 100 U/mL penicillin and 100 μg/mL streptomycin supplemented with 20 ng/mL M-CSF (BioLegend) at a density of 2×10^5^ cells/mL. Cells were incubated at 37 ^°^C, 5% CO_2_, and the medium was exchanged on day 3 and 5 of the culture. On day 7, the BMDMs were stimulated with 10 μM LPA for 4 hours and collected in TRIzol for RNA extraction.

### Transwell migration assay

5×10^5^ BMDMs were seeded on inserts with 5 μm pore size (Corning). RPMI 1640 containing 2% FBS with 10 μM LPA was placed in the outer chamber. The cells were allowed to migrate for 18 hours. Migrated cells and the cells in the upper chamber were collected and resuspended in 200 μl of FACS buffer. 70 μl of each sample was acquired using BD Accuri^TM^ C6 flow cytometer (BD Biosciences) and percentage of migrated cells was calculated.

### Enzyme linked immunosorbent assays (ELISA) for corticosterone detection

Corticosterone concentrations were determined in mouse colonic explants, which had been incubated for 24 hours in 24-well plates in 500 μl of DMEM containing 2% FBS and 100 U/mL penicillin and 0.1 mg/mL streptomycin. Corticosterone levels were determined using the Corticosterone Competitive ELISA kit (Invitrogen) and normalized to the weights of colon pieces measured before the assay.

### Statistical analysis

Data are presented as dot plots of individual values with medians. GraphPad Prism software was used to graph the data and calculate statistical significance. P values were calculated using either unpaired t-test, Mann-Whitney U or two-way ANOVA tests depending on the experimental setting. Data were further analyzed by Grubbs’ test to identify the outliers. For all tests p values were indicated as followed: *: p≤0.05, **: p≤0.005, ***: p≤0.0005

## Supplemental Information

Supplemental Information includes a list of the Swiss IBD Cohort Investigators members and affiliations, 7 figures, and 3 tables.

## Supplemental Figure Legends

**Figure S1. Zebrafish, Mouse, and Human GPR35 Proteins**

(A) ClustalW alignment of mouse and human GPR35 with Gpr35a and Gpr35b paralogs identified in zebrafish (red). Alignment scores per pair of sequences were calculated by ClustalW.

(B) Phylogenetic tree including protein sequences from human, mouse, and zebrafish *GPR35* and *GPR55* orthologs. Analysis was performed by ClustalW.

(C) *gpr35a* and *gpr35b* mRNA levels in the dissected intestine or rest of the body from WT zebrafish larvae. Target genes were normalized to *ef1a* housekeeping gene. One representative experiment is shown from two experiments.

(D) *GPR35* mRNA levels retrieved from the Human Protein Atlas (https://www.proteinatlas.org).

**Figure S2. *Gpr35* Is Expressed in Colon Tissue in Mice**

(A) Construct design of *Gpr35*-tdTomato reporter mice.

(B) Immunofluorescence staining of GPR35 and secondary antibody control in small intestine from *Gpr35*-tdTomato mice. Sections were stained for GPR35 (red) and NucBlue (blue) for nuclear staining. Scale bars, 50 μm.

(C) Percentage of *Gpr35*-tdTomato-positive B cells, CD4^+^ T cells, CD8^+^ T cells, neutrophils, dendritic cells, NK cells, ILC1 cells, ILC2 cells, and ILC3 cells in the colonic lamina propria of *Gpr35*-tdTomato reporter mice (red unfilled histograms) and WT mice (gray histograms) as the background control.

(D) Gating strategy for the analysis of colonic lamina propria monocytes and macrophages. Lin: lineage to exclude CD3, CD19, and NK1.1^+^ cells.

(E) Unsupervised heatmap expression profile from RNA sequencing of *Gpr35*-tdTomato-positive (Pos) and -negative (Neg) colonic lamina propria macrophages (co-LP MAC).

**Figure S3. Construction of *gpr35b^uu1892^* Mutant Zebrafish**

(A) Schematic representation of the *gpr35b* (ENSDARG00000075877) locus. The *gpr35b^uu1892^* mutant line was generated by deletion of a 10-bp fragment within exon 2 (blue box).

(B) Schematic of the resulting Gpr35b proteins from WT or *gpr35b^uu19b2^* mutant fish. Deletion results in a preliminary stop codon after the first 27 amino acids.

**Figure S4. Design of *Gpr35*^-/-^ Mice**

(A) Construct design for production of *Gpr35*^-/-^ mice.

(B) Immunofluorescence staining of GPR35 in colon from *Gpr35*^-/-^ (right) and WT mice (left). Sections were stained for GPR35 (red) and NucBlue (blue) for nuclear staining. Scale bars, 50 μm.

**Figure S5. *Gpr35*-Deficient Mice Show Exacerbated DSS Colitis**

(A) Percentages of body weight (normalized to initial weight) of untreated (UT) or DSS-treated WT or *Gpr35*^-/-^ mice. Data are shown as mean ± SD for four mice per group.

(B) Disease activity scores assessed daily by monitoring UT or DSS-treated WT or *Gpr35*^-/-^ mice. Data are shown as mean ± SD for four mice per group.

(C) Representative images of colons from UT or DSS-treated WT or *Gpr35*^-/-^ mice on day 8.

(D) Colon lengths measured from colon images as shown in (C) of UT or DSS-treated WT or *Gpr35*^-/-^ mice on day 8.

(E) H&E staining of colon tissue sections from UT or DSS-treated WT or *Gpr35*^-/-^ mice taken on day 8. Scale bars, 100 μm.

(F) Histology scores obtained from H&E staining of colons as shown in E.

Data are represented as individual values, with each dot representing one mouse with medians (C-F). *p < 0.05, **p < 0.01, *** < p 0.001 by two-way ANOVA with Tukey’s multiple comparisons test (A, B) or Mann-Whitney (D, F).

**Figure S6. Design of Conditional *Gpr35*^flox^ Mice**

(A) Construct design for production of *Gpr35*^flox^ mice.

(B) After mating of *Gpr35*^flox^ with *Cx3cr1^CreER^* to obtain *Gpr35^ΔCx3cr1^*mice, colon was taken from *Gpr35^ΔCx3cr1^*mice injected i.p. with vehicle (left) or tamoxifen (right). Sections were stained for GPR35 (red) and NucBlue (blue) for nuclear staining. Arrowheads indicate CX3CR1-YFP^+^ macrophages. Scale bars, 50 μm.

(C) Percentages of GPR35^+^ cells among CX3CR1-YFP^+^ cells. Data are represented as individual values, with each dot representing one biological replicate with medians. *p ≤ 0.05 by Mann-Whitney.

**Figure S7. DSS-treated *Gpr35^ΔCx3cr1^* mice Have Increased Neutrophil Numbers**

(A-C) Flow cytometric analysis of colonic lamina propria (co-LP) macrophages (MAC), dendritic cells (DCs), neutrophils and monocyte subsets (Ly6C^high^ to Ly6C^low^) of vehicle-treated (corn oil) or tamoxifen (TMX)-treated WT or *Gpr35^ΔCx3cr1^* mice on day 7 of DSS colitis. Quantification of flow cytometric data (B) for frequency (C) and number of macrophages, DCs, Ly6^hi^, Ly6^mid^ or Ly6 ^low^ monocytes and neutrophils is shown below.

(D) mRNA expression levels of colonic *Il10*, *Il1b, Il6*, and *Tnf* relative to *Actb* by qRT-PCR of vehicle- or TMX-treated WT or *Gpr35^ΔCx3cr1^* mice on day 7 of DSS colitis. Data are presented as individual values with each dot representing one animal with medians. *p ≤ 0.05, **p ≤ 0.01, ***p ≤0.001 by Mann-Whitney.

## Supplemental Tables

**Table S1.**
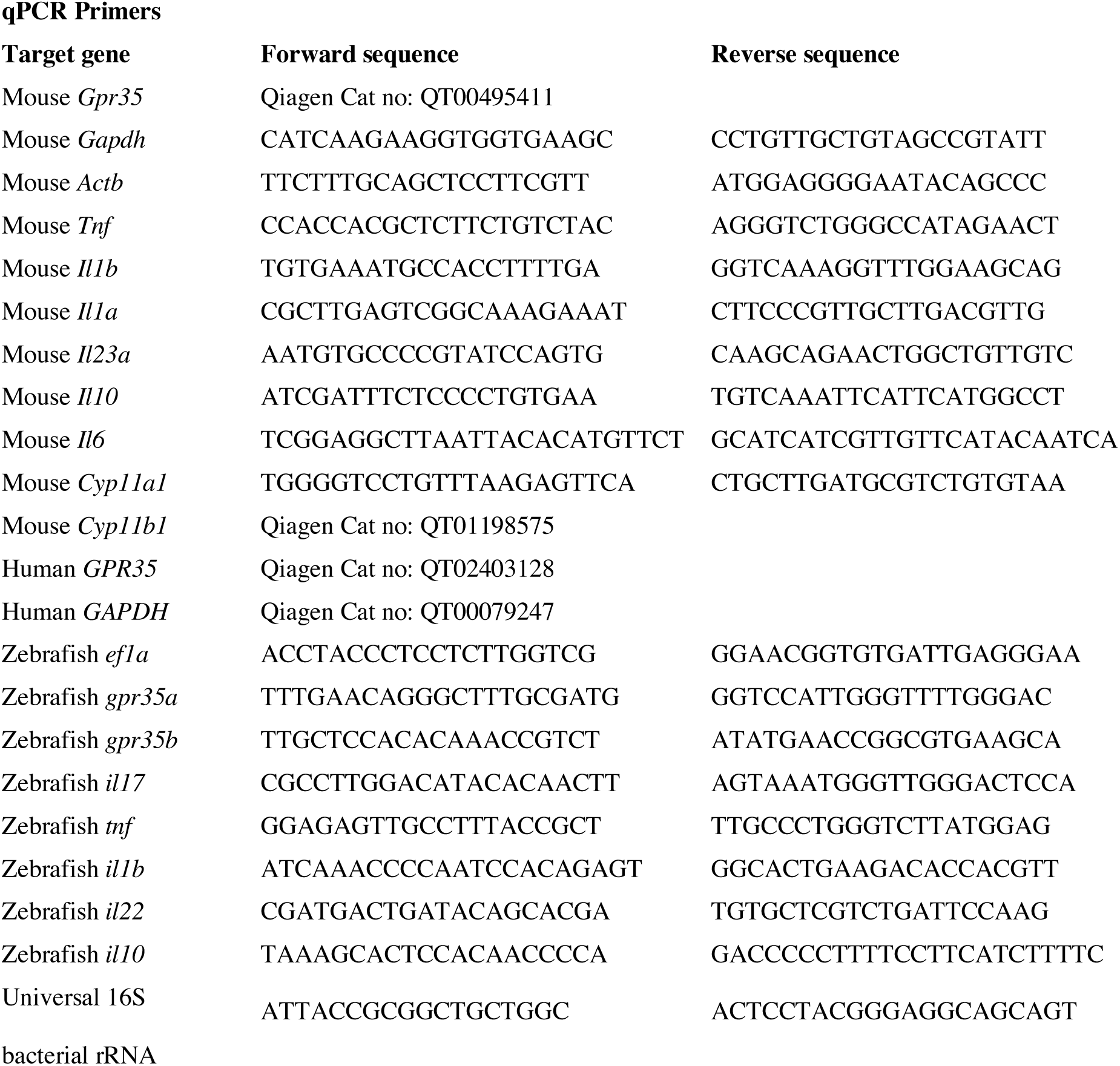
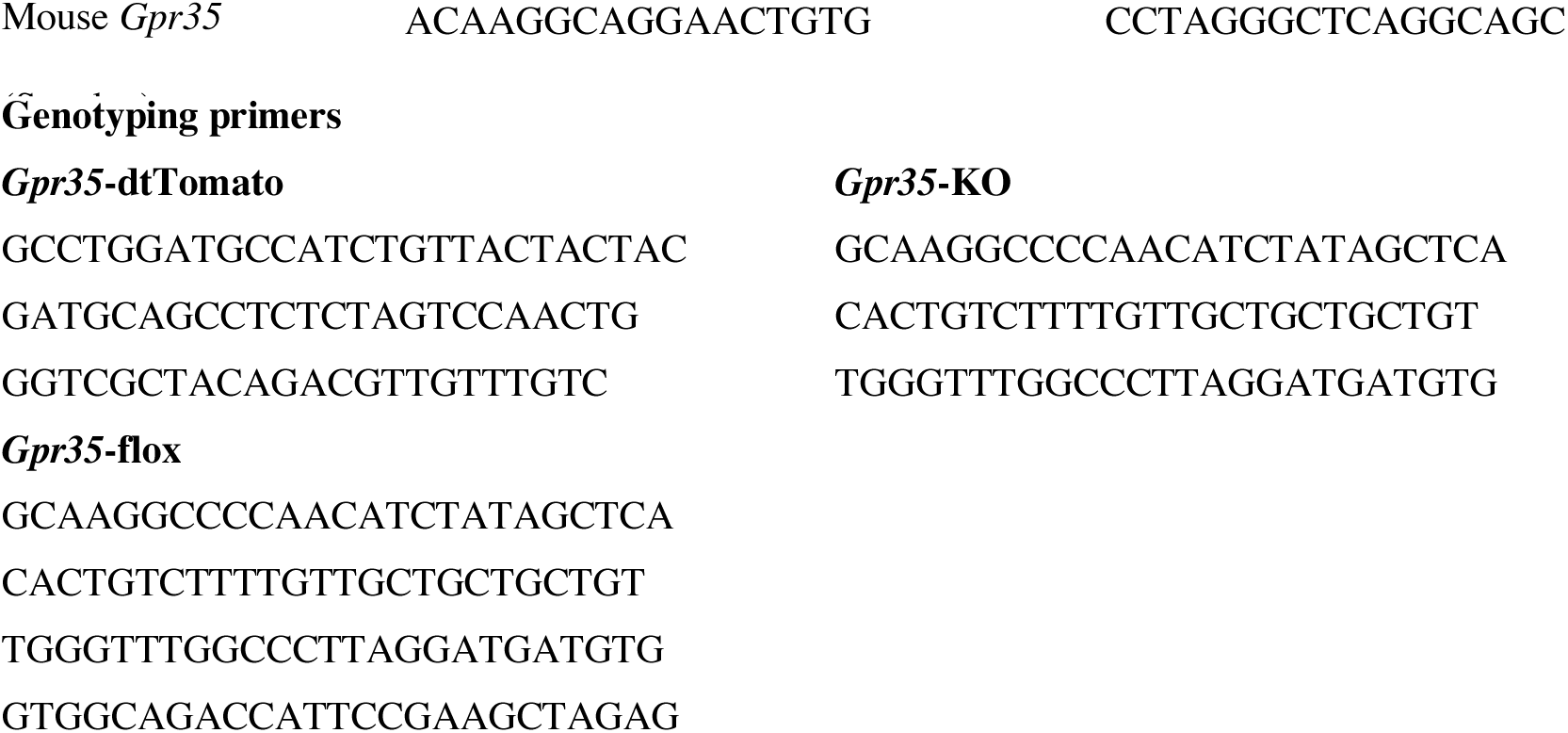
Primers.

**Table S2.** Disease Characteristics of Swiss IBD Cohort Study Group Patients Who Provided Biopsies for Expression Analysis.

Baseline Group Characteristics
|  | Crohn's disease |  | Ulcerative colitis |  |
| --- | --- | --- | --- | --- |
|  | Quiescent (n=20) | Active (n=11) | Quiescent (n=20) | Active (n=11) |
| Gender (male/female %, n) | 60/40 %<br>12 male, 8 female | 36/64 %<br>4 m, 7 f | 35/65%<br>12 m, 8 f | 54/45%<br>6 m, 5 f |
| Median age (range), yr | 56 (32-81) | 39 (31-71) | 58 (31-81) | 53 (24-68) |
| Mean BMI (SD), kg/m<sup>2</sup> | 24.15 (3) | 25.55 (4.9) | 24.70 (2.99) | 23.89 (4.08) |
| Median age at diagnosis (range), yr | 33.5 (18-60) | 21 (14-34) | 28.5 (14-65) | 30 (16-60) |
| Median disease duration (range), yr | 21.5 (6-45) | 19 (7-37) | 26 (8-44) | 16 (7-38) |
| **CD extent, n (%)** |  |  |  |  |
| Ileum isolated | 5 (25%) | 2 (18%) |  |  |
| Colon isolated | 2 (10%) | 3 (27%) |  |  |
| Ileocolonic | 2 (10%) | 5 (46%) |  |  |
| Unknown | 11 (55%) | 1 (9%) |  |  |
| **UC extent, n (%)** |  |  |  |  |
| Proctitis |  |  | 1 (5%) | 1 (10%) |
| Left-sided colitis |  |  | 3 (20%) | 5 (45%) |
| Pancolitis |  |  | 5 (25%) | 5 (45%) |
| Unknown |  |  |  |  |
| **Current medical treatment, n (%)** |  |  |  |  |
| No treatment | 5 (25%) |  | 5 (25%) | 1 (9%) |
| 5-ASA | 6 (30%) |  | 13 (65%) | 9 (81%) |
| Steroids |  | 6 (54%) |  | 4 (36%) |
| Immunosuppressants | 11 (55%) | 1 (9%) | 4 (20%) | 1 (9%) |
| Anti-TNF |  | 1 (9%) |  | 3 (27%) |
| **Smoking status** |  |  |  |  |
| Non-smoker, n (%) | 12 (60%) | 6 (54%) | 15 (75%) | 9 (81%) |
| Active smoker, n (%) | 6 (30%) | 5 (46%) | 1 (5%) | 1 (9.5%) |
| Unknown | 2 (10%) | 0 | 4 (20%) | 1 (9.5%) |

**Table S3.** Characteristics of Basel IBD patients Who Provided Biopsies for Immunofluorescence.

| Patient ID | Gender | Age | BMI | Age at diagnosis | Smoking status | Location inflamed | Location non-inflamed | Treatment at time of study | DAI |
| --- | --- | --- | --- | --- | --- | --- | --- | --- | --- |
| **Ulcerative colitis** |  |  |  |  |  |  |  |  |  |
| 504 | F | 50 | 25.6 | 34 | Unk. | sigma/rectum | trans. colon | none | 5 |
| 535 | F | 71 | 27.3 | 56 | Unk. | rectum/sigmoid | colon | none | 6 |
| 551 | F | 53 | 25 | 37 | Unk. | rectum | Unk. | Unk. | Unk. |
| 619 | Unk. | Unk. | Unk. | Unk. | Unk. | sigmoid/rectum | ascend. / trans. colon | None, Salofalk 10 days before | Unk. |
| **Crohn's disease** |  |  |  |  |  |  |  |  |  |
| 558 | F | 72 | 19 | 23 | non-smoker | Unk | rectum | Quantalan, Immodium | Unk. |
| 568 | M | 68 | 32 | 52 | active | sigmoid | desc. colon | Spiricort, Aldactone, Orfiril | 74 |
| 620 | F | 69 | 21.6 | 56 | active | term. Ileum | ascend. colon | Unk | 70 |

Unk, unknown; DAI, disease activity index

## Swiss IBD Cohort Investigators

Karim Abdelrahman^6^, Gentiana Ademi^7^, Patrick Aepli^8^, Claudia Anderegg^9^, Anca-Teodora Antonino^10^, Eva Archanioti^31^, Eviano Arrigoni^11^, Diana Bakker de Jong^3^, Bruno Balsiger^12^, Polat Bastürk^3^, Peter Bauerfeind^13^, Andrea Becocci^14^, Dominique Belli^14^, José M. Bengoa^11^, Luc Biedermann^15^, Janek Binek^16^, Mirjam Blattmann^15^, Stephan Boehm^17^, Tujana Boldanova^3^, Jan Borovicka^7^, Christian P. Braegger^19^, Stephan Brand^7^, Lukas Brügger^20^, Simon Brunner^3^, Patrick Bühr^19^, Sabine Burk^15^, Bernard Burnand^21^, Emanuel Burri^22^, Sophie Buyse^23^, Dahlia-Thao Cao^24^, Ove Carstens^20^, Dominique H. Criblez^8^, Sophie Cunningham^11^, Fabrizia D’Angelo^25^, Philippe de Saussure^11^, Lukas Degen^3^, Joakim Delarive^26^, Christopher Doerig^27^, Barbara Dora^15^, Susan Drerup^28^, Mara Egger^21^, Ali El-Wafa^29^, Matthias Engelmann^30^, Jessica Ezri^31^, Christian Felley^32^, Markus Fliegner^33^, Nicolas Fournier^21^, Montserrat Fraga^31^, Yannick Franc^21^, Remus Frei^7^, Pascal Frei^34^, Michael Fried^15^, Florian Froehlich^35^, Raoul Ivano Furlano^36^, Luca Garzoni^37^, Martin Geyer^38^, Laurent Girard^11^, Marc Girardin^39^, Delphine Golay^21^, Ignaz Good^40^, Ulrike Graf Bigler^20^, Beat Gysi^41^, Johannes Haarer^7^, Marcel Halama^42^, Janine Haldemann^12^, Pius Heer^43^, Benjamin Heimgartner^20^, Beat Helbling^34^, Peter Hengstler^16^, Denise Herzog^44^, Cyrill Hess^8^, Roxane Hessler^27^, Klaas Heyland^45^, Thomas Hinterleitner^46^, Claudia Hirschi^30^, Petr Hruz^3^, Pascal Juillerat^20^, Stephan Kayser^47^, Céline Keller^48^, Carolina Khalid-de Bakker^3^, Christina Knellwolf(-Grieger)^7^, Christoph Knoblauch^49^, Henrik Köhler^9^, Rebekka Koller^19^, Claudia Krieger(-Grübel)^7^, Patrizia Künzler^7^, Rachel Kusche^9^, Frank Serge Lehmann^43^, Andrew J. Macpherson^20^, Michel H. Maillard^31,48^, Michael Manz^3^, Astrid Marot^21^, Rémy Meier^50^, Christa Meyenberger^7^, Pamela Meyer^7^, Pierre Michetti^31,48^, Benjamin Misselwitz^15^, Patrick Mosler^51^, Christian Mottet^52^, Christoph Müller^53^, Beat Müllhaupt^15^, Leilla Musso^21^, Michaela Neagu^54^, Cristina Nichita^55^, Jan H. Niess^3^, Andreas Nydegger^31,48^, Nicole Obialo^15^, Diana Ollo^32^, Cassandra Oropesa^32^, Ulrich Peter^45^, Daniel Peternac^56^, Laetitia Marie Petit^32^, Valérie Pittet^21^, Daniel Pohl^15^, Marc Porzner^57^, Claudia Preissler^58^, Nadia Raschle^15^, Ronald Rentsch^59^, Sophie Restellini^32^, Alexandre Restellini^39^, Jean-Pierre Richterich^8^, Frederic Ris^32^, Branislav Risti^60^, Marc Alain Ritz^61^, Gerhard Rogler^15^, Nina Röhrich^7^, Jean-Benoît Rossel^21^, Vanessa Rueger^19^, Monica Rusticeanu^20^, Markus Sagmeister^63^, Gaby Saner^12^, Bernhard Sauter^64^, Mikael Sawatzki^7^, Michael Scharl^15^, Martin Schelling^7^, Susanne Schibli^65^, Hugo Schlauri^66^, Dominique Schluckebier^32^, Sybille Schmid(-Uebelhart)^20^, Daniela Schmid^49^, Jean-François Schnegg^67^, Alain Schoepfer^31,48^, Vivianne Seematter^21^, Frank Seibold^12^, Mariam Seirafi^68^, Gian-Marco Semadeni^7^, Arne Senning^19^, Christiane Sokollik^65^, Joachim Sommer^21^, Johannes Spalinger^8,65^, Holger Spangenberger^69^, Philippe Stadler^70^, Peter Staub^71^, Dominic Staudenmann^8^, Volker Stenz^72^, Michael Steuerwald^61^, Alex Straumann^43^, Bruno Strebel^20^, Andreas Stulz^8^, Michael Sulz^7^, Aurora Tatu^20^, Michela Tempia-Caliera^73^, Amman Thomas^62^, Joël Thorens^74^, Kaspar Truninger^75^, Radu Tutuian^20^, Patrick Urfer^76^, Stephan Vavricka^15^, Francesco Viani^77^, Jürg Vögtlin^61^, Roland Von Känel^15^, Dominique Vouillamoz^78^, Rachel Vulliamy^21^, Paul Wiesel^55^, Reiner Wiest^20^, Stefanie Wöhrle^8^, Tina Wylie^79^, Samuel Zamora^39^, Silvan Zander^80^, Jonas Zeitz^15^, Dorothee Zimmermann^76^ Clinique de Montchoisi, Lausanne, Switzerland; ^7^Kantonsspital St. Gallen, St. Gallen, Switzerland; ^8^Kantonsspital Luzern, Luzern, Switzerland; ^9^Kantonspital Aarau, Klinik für Kinder und Jugendliche, Aarau, Switzerland; ^10^Hôpital Riviera–Site du Samaritain, Vevey, Vaud, Switzerland; ^11^GI private practice, Geneva, Switzerland; ^12^Gastroenterologische Praxis, Bern, Switzerland; ^13^Department Gastroenterology and Hepatology, Stadtspital Triemli, Zurich, Switzerland; ^14^Department of Pediatric, Geneva University Hospital, Geneva, Switzerland; ^15^Department of Gastroenterology and Hepatology, University Hospital Zurich, University of Zurich, Zurich, Switzerland; ^16^Gastroenterologie am Rosenberg, St. Gallen, Switzerland; ^17^Spital Bülach, Bülach, Zurich, Switzerland; ^18^Department of Biomedicine, University of Basel, Basel, Switzerland; ^19^University Children’s Hospital, Zurich, Switzerland; ^20^Department of Visceral Surgery and Medicine, Bern University Hospital, University of Bern, Bern, Switzerland; ^21^Institute of Social and Preventive Medicine (IUMSP), Lausanne University Hospital, Lausanne, Switzerland; ^22^Department Gastroenterology, Kantonsspital Liestal, Liestal, Switzerland; ^23^GI private practice, Yverdon-les-Bains, Switzerland; ^24^Hôpital Neuchâtelois, La Chaux-de-fonds, Neuchâtel, Switzerland; ^25^Department Gastroenterology and Hepatology, Geneva University Hospital, Geneva, Switzerland; ^26^GI private practice, Lausanne, Switzerland; ^27^Clinique Cecil, Lausanne, Switzerland; ^28^Schulthess Clinic, Zurich, Switzerland; ^29^GI private practice, La Chaux-de-Fonds, Switzerland; ^30^Gastropraxis Luzern, Luzern, Switzerland; ^31^Service of Gastroenterology and Hepatology, Department of Medicine, Centre HospitalierUniversitaire Vaudois and University of Lausanne, Lausanne, Switzerland; ^32^Centre de Gastroentérologie Beaulieu SA, Geneva, Switzerland; ^33^Medical Center Sihlcity, Zurich, Switzerland; ^34^Gastroenterologie Bethanien, Zurich, Switzerland; ^35^Hospital of the Canton of Jura, Porrentruy And Delémont, Jura, Switzerland; ^36^Universitäts-Kinderspital beider Basel (UKBB), Basel, Switzerland; ^37^Clinique des Grangettes, Geneva University Hospital, Genève, Switzerland; ^38^GI private practice, Wettingen, Aargau, Switzerland; ^39^Groupe Médical d’Onex, Onex, Switzerland; ^40^Spital Walenstadt, Walenstadt, St. Gallen, Switzerland; ^41^GI private practice, Reinach, Switzerland; ^42^Aerztehaus Fluntern, Zurich, Switzerland; ^43^GI private practice, Olten, Switzerland; ^44^HFR Hôpital fribourgeois–Pédiatrie, Fribourg, Switzerland; ^45^KSW Kantonsspital Winterthur Kinderklinik, Winterthur, Switzerland; ^46^GI private practice, Zurich, Switzerland; ^47^GI private practice, Luzern, Switzerland; ^48^Gastroenterology La Source-Beaulieu, Lausanne, Switzerland; ^49^Kantonsspital Nidwalden, Stans, Nidwalden, Switzerland; ^50^AMB – Arztpraxis MagenDarm Basel, Basel, Switzerland; ^51^Kantonsspital Graubünden, Chur, Switzerland; ^52^GI private practice, Sion, Switzerland; ^53^Division of Experimental Pathology, Institute of Pathology, University of Bern, Bern, Switzerland; ^54^Spital Tiefenau, Bern, Switzerland; ^55^Centre médical d’Epalinges, Epalinges, Switzerland; ^56^Spital Waid, Zurich, Switzerland; ^57^Spital Lachen, Lachen, Switzerland; ^58^Kantonsspital Olten, Olten, Switzerland; ^59^GI private practice, St. Gallen, Switzerland; ^60^GI practice, Dietikon, Switzerland; ^61^GI practice, Liestal, Switzerland; ^62^GI private practice, Waldkirch, St. Gallen, Switzerland; ^63^GI private practice, Heerbrugg, Switzerland; ^64^Klinik Hirslanden Zürich, Zurich, Switzerland; ^65^Kinderklinik Bern, Bern University Hospital, Bern, Switzerland; ^66^Derby Center, Wil, Switzerland; ^67^GI private practice, Montreux, Switzerland; ^68^Clinique La Colline, Geneva, Switzerland; ^69^Kantonsspital Wolhusen, Wolhusen, Switzerland; ^70^GI private practice, Payerne, Switzerland; ^71^Spital Heiden Appenzell Ausserrhoden, Heiden, Switzerland; ^72^Kantonsspital Münsterlingen, Münsterlingen, Switzerland; ^73^Clinique des Grangettes, Chêne-Bougeries, Switzerland; ^74^GI private practice, Yverdon, Switzerland; ^75^GI private practice, Langenthal, Switzerland; ^76^Hirslanden Klinik Aarau, Gastro Zentrum, Aarau, Switzerland; ^77^Private practice, Vevey, Switzerland; ^78^Private practice, Pully, Switzerland; ^79^Infirmière de Recherche chez CHUV Lausanne University Hospital, Lausanne, Switzerland; ^80^Spital Limmattal, Schlieren, Switzerland

