## Supplementary Figures for "Lysophosphatidic acid-mediated GPR35 signaling in CX3CR1^+^ macrophages regulates the intestinal cytokine milieu"

### Figure S1

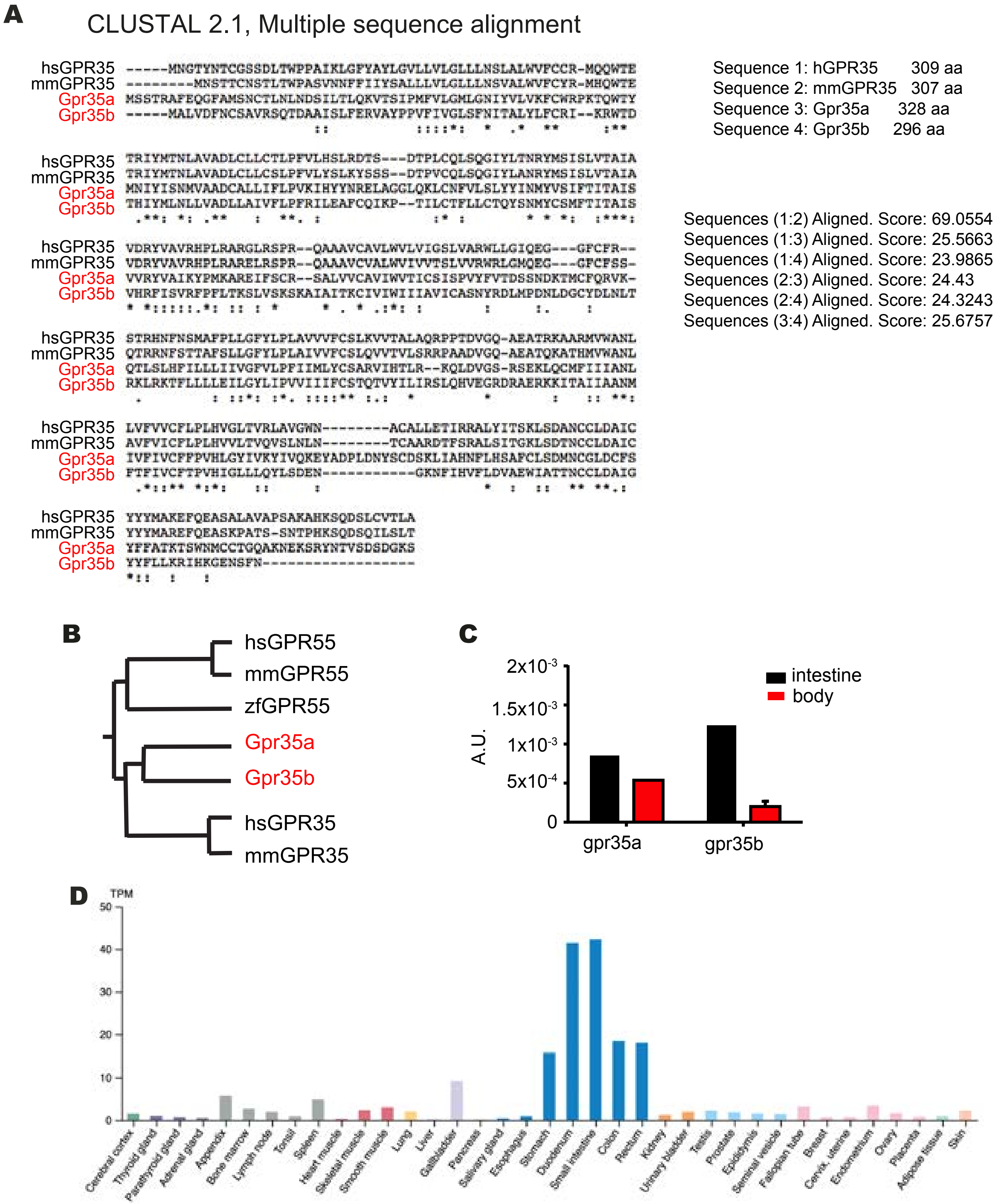


**Figure S1. Zebrafish GPR35 proteins**

(A) ClustalW alignment of mouse and human GPR35 with Gpr35a and Gpr35b paralogs identified in zebrafish (red fonts). Alignment scores per pair of sequences were calculated by ClustalW.
(B) Phylogenetic tree including protein sequences from human, mouse and zebrafish *GPR35 and GPR55* orthologs. The analysis was made by ClustalW.
(C) *gpr35a* and *gpr35b* mRNA levels in the dissected intestine or rest of the body from WT zebrafish embryos. Target genes were normalized to *ef1a* housekeeping gene. One representative experiment out of two.
(D) *Gpr35* mRNA levels retrieved from the Human Protein Atlas (https://www.proteinatlas.org)

### Figure S2

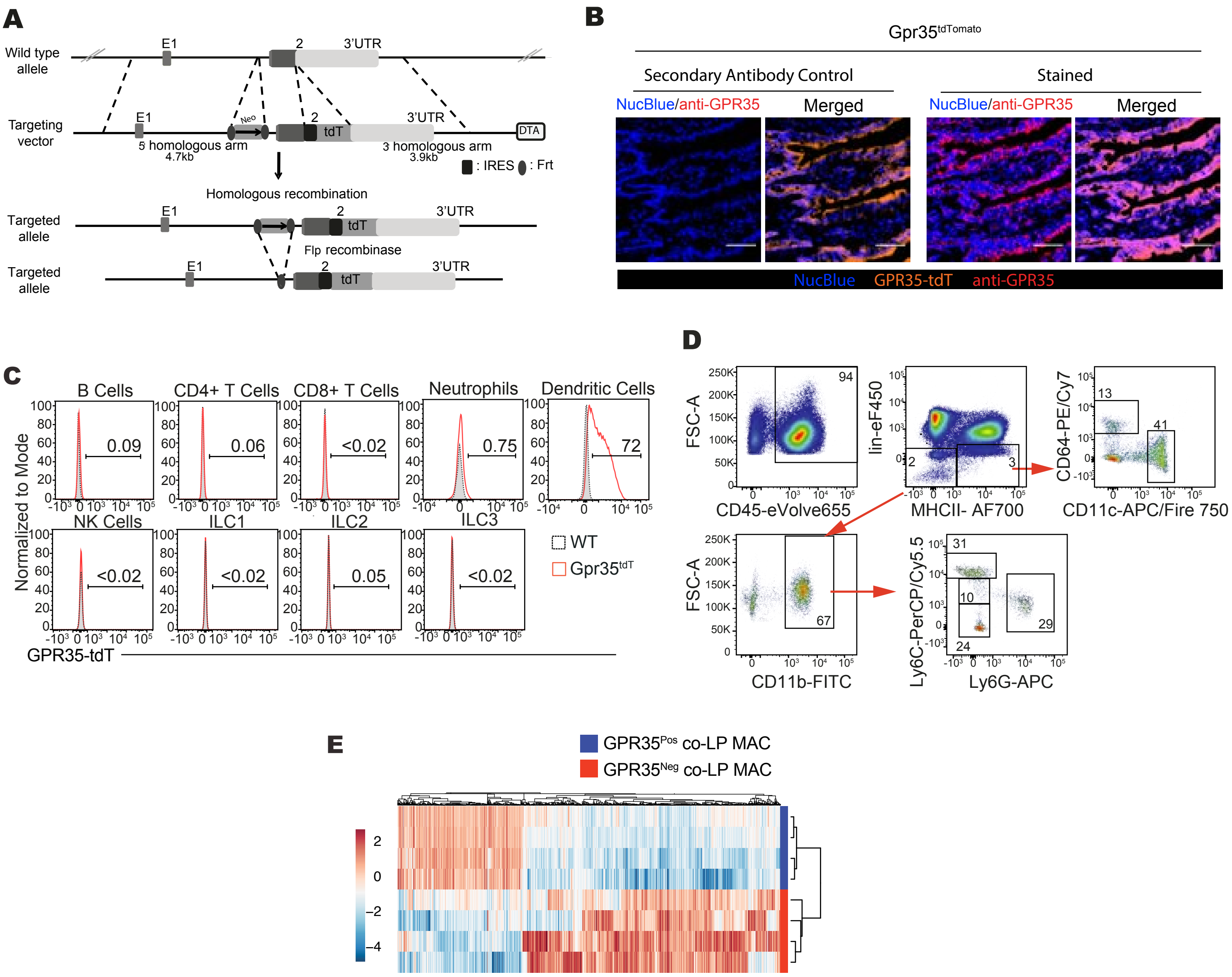


**Figure S2. *Gpr35* is expressed in the colon.**
(A) Construct design of *Gpr35*-tdTomato reporter mice.
(B) Immunofluorescence staining of GPR35 on small intestine taken from *Gpr35*-tdTomato mice (right) and secondary antibody control (left). Sections were stained for GPR35 (red) and NucBlue (blue) for nuclear staining. Scale bars represent 50 μm.
(C) Gpr35-tdTomato⁺ B, CD4 T, CD8 T, dendritic and NK cells, neutrophils, ILC1, ILC 2 and ILC 3 from the colonic lamina propria of *Gpr35*-tdTomato reporter mice (blank red histograms) and wt mice (gray histograms) as the background control.
(D) Gating strategy for the analysis of colonic lamina propria monocytes and macrophages. Lin: lineage to exclude CD3, CD19 and NK1.1⁺ cells.
(E) Unsupervised heatmap expression profile from RNA sequencing of GPR35-tdTomato positive (Pos) and negative (Neg) colonic lamina propria macrophages (co-LP MAC).
Data are represented as individual values with medians. *p 0.05, **p 0.01, ***p 0.001 by two-way ANOVA with Tukey's multiple comparisons test.

### Figure S3

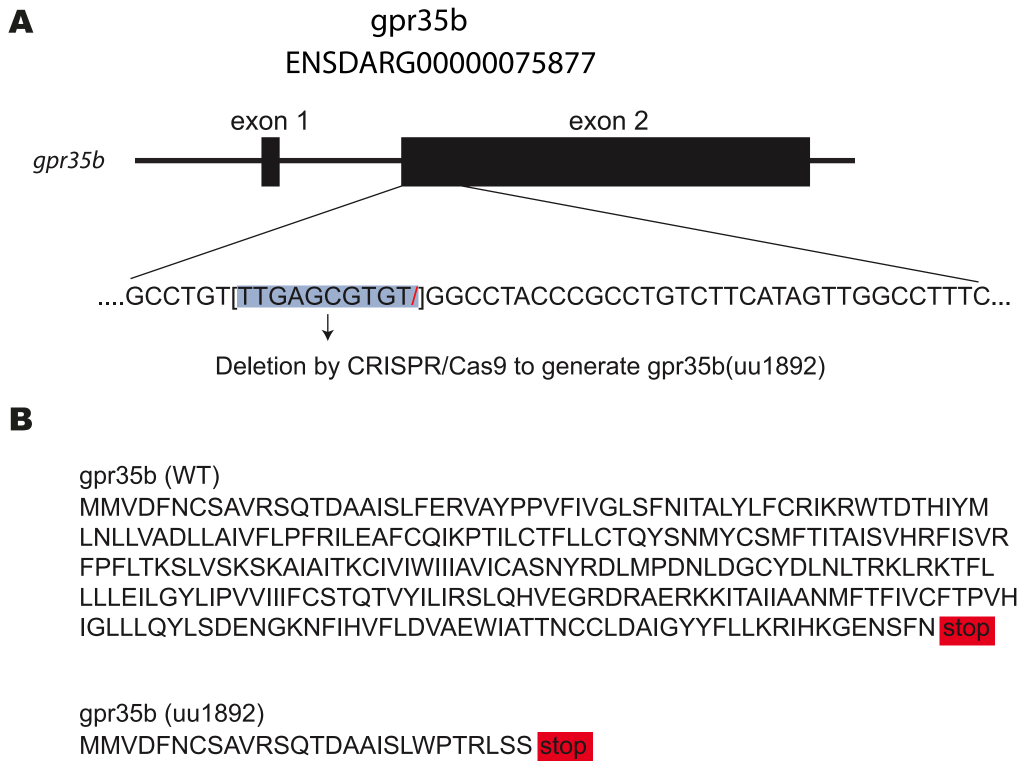


**Figure S3. Construction of *gpr35b$^{uu1892}$* mutant zebrafish**

(A) Schematic representation of the *gpr35b* (ENSDARG00000075877) locus. The *gpr35b$^{uu1892}$* mutant line was generated by deletion of a 10bp fragment within exon 2 (blue box).

(B) Schematic representation of the resulting Gpr35b proteins from WT or *gpr35b$^{uu1892}$*. Deletion results in a preliminary stop codon after the first 27 aminoacids.

### Figure S4

**A**

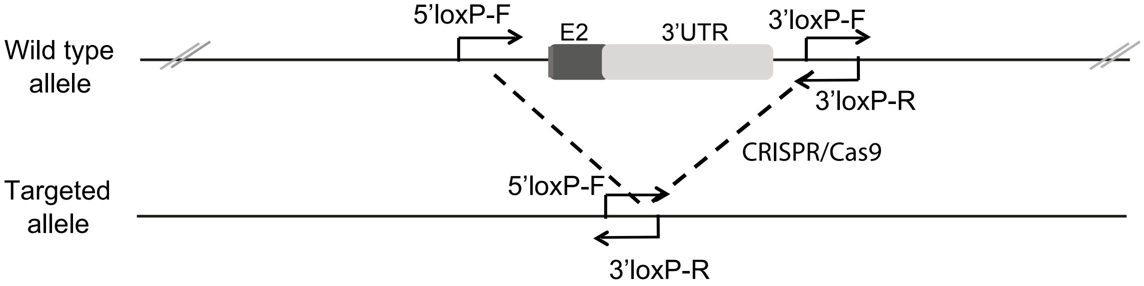


**B**

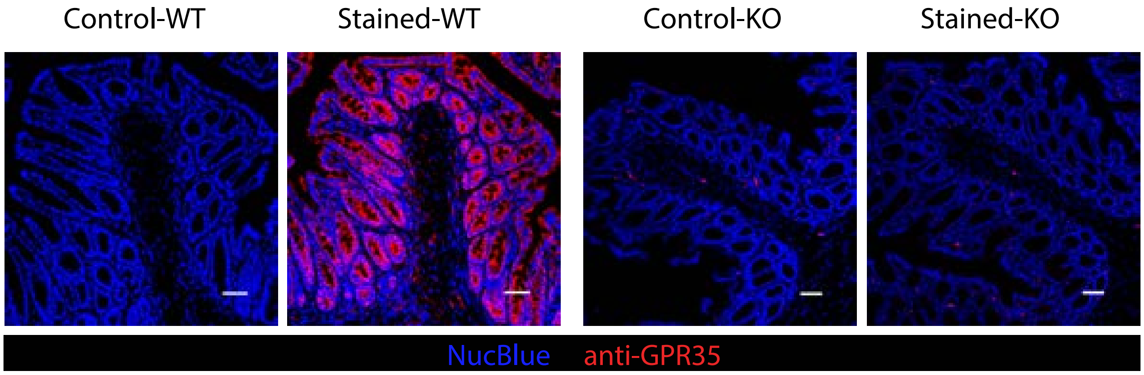


**Figure S4. Design of global GPR35$^{-/-}$ m ice**
(A) Construct design of *Gpr35*-KO mice.
(B) Immunofluorescence staining of GPR35 on colon taken from *Gpr35*-KO (right) and wt mice (left). Sections were stained for GPR35 (red) and NucBlue (blue) for nuclear staining. Scale bars represent 50 µm.

### Figure S5

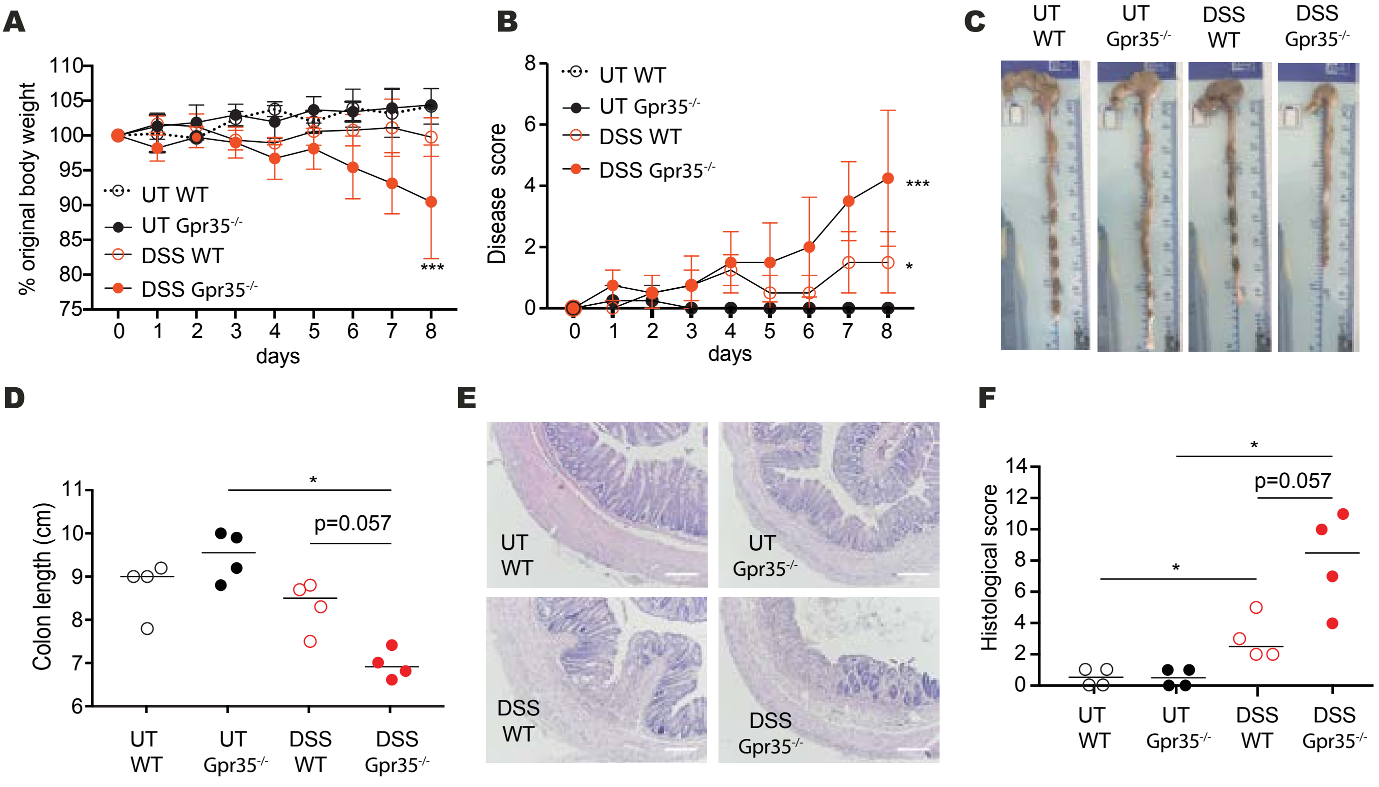


**Figure S5. Increased DSS colitis in GPR35-deficient mice.**

(A) Percentages of body weights for 8 days compared to initial weight of untreated (UT) or DSS-treated wt or *Gpr35*$^{-/-}$ mice. Data are shown as mean ± SD for 4 individual mice per group.

(B) Disease activity scores assessed daily for 8 days by monitoring untreated (UT) or DSS-treated wt or Gpr35$^{-/-}$ mice. Data are shown as mean ± SD for 4 individual mice per group.

(C) Representative images of colons from untreated (UT) or DSS-treated wt or Gpr35$^{-/-}$ mice on day 8.

(D) Colon lengths measured from colon images (C) of UT or DSS-treated wt or Gpr35$^{-/-}$ mice on day 8.

(E) H&E staining of colon tissue sections from untreated (UT) or DSS-treated wt or Gpr35$^{-/-}$ mice taken on day 8. Scale bars represent 100 µm.

(F) Histology scores obtained from H&E staining of colons.

Data are represented as individual values with medians (C-F). *p≤0.05, **p≤ 0.01, ***p ≤0.001, ****p≤0.0001 by two-way ANOVA with Tukey's multiple comparisons test (A, B) or Mann-Whitney (D, F).

### Figure S6

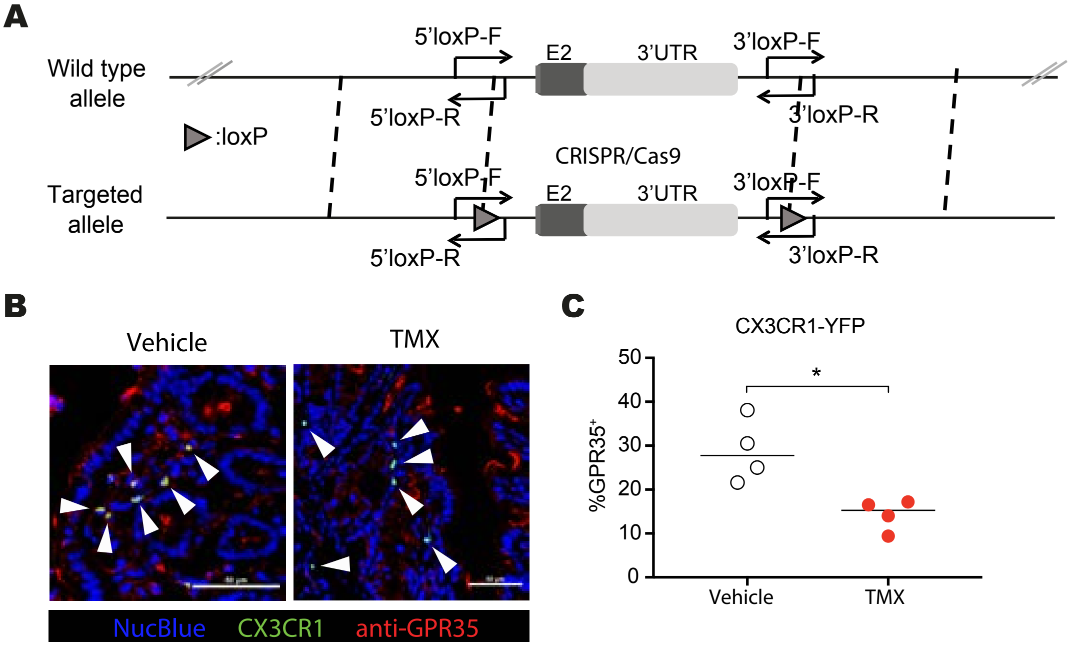


**Figure S6. Design and validation of conditional GPR35$^{-/-}$ mice**
(A) Construct design of *Gpr35*-flox mice.
(B) Immunofluorescence staining of GPR35 on colon taken from *Gpr35*$^{\Delta Cx3cr1}$ mice injected i.p. with vehicle (left) or tamoxifen (right). Sections were stained for GPR35 (red) and NucBlue (blue) for nuclear staining. White arrowheads indicate macrophages. Scale bars represent 50 µm.
(C) Percentages of GPR35$^{+}$ cells among CX3CR1-YFP$^{+}$ cells. Data are represented as individual values with medians (C-F). $^{*}p \leq 0.05$ by Mann-Whitney.
!

### Figure S7

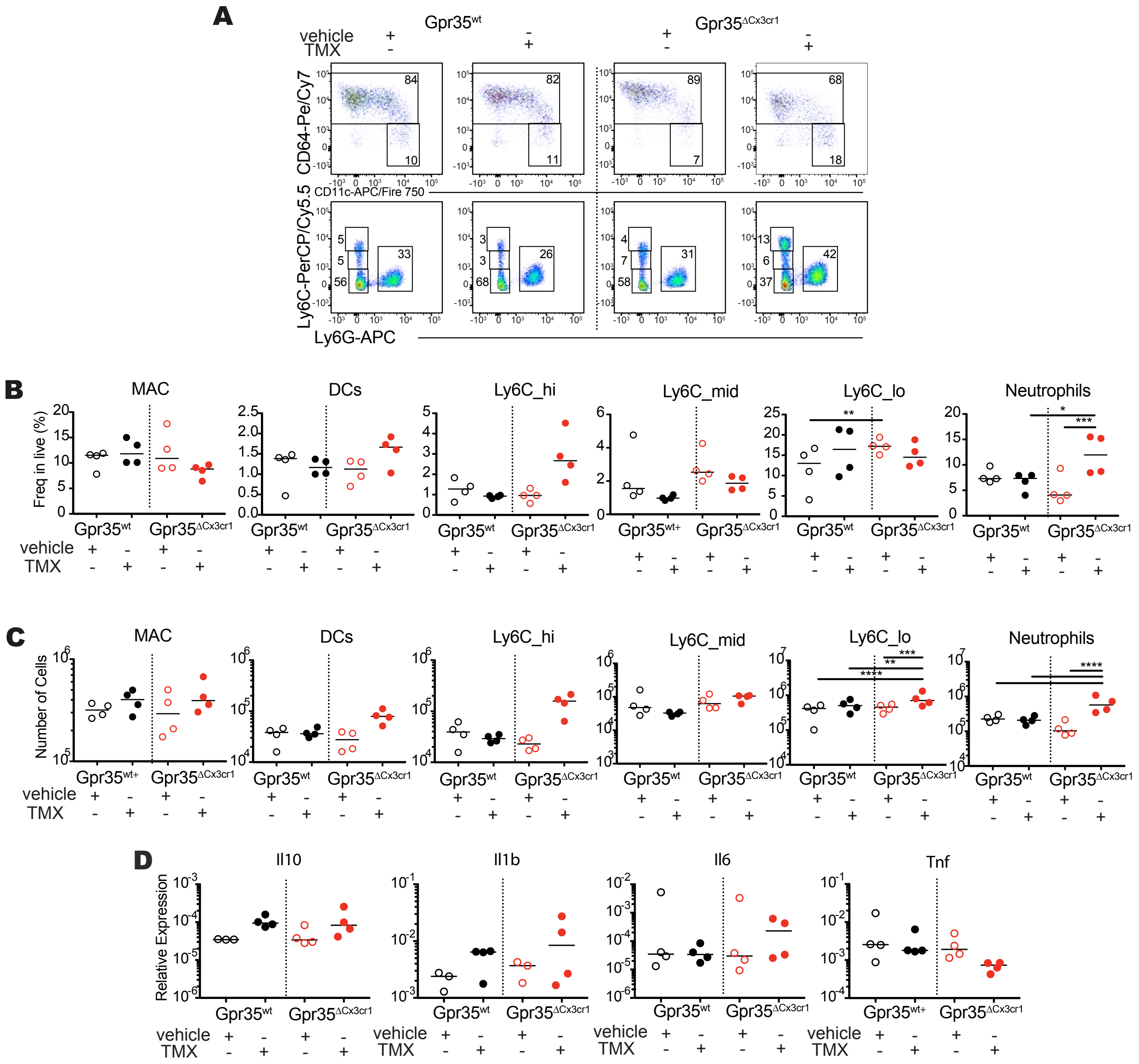


**Figure S7. Increased neutrophil num bers in DSS-treated *Gpr35$^{\Delta Cx3cr1}$* mice.**
(A) Flow cytometric analysis of colonic lamina propria (co-LP) macrophages (MAC), dendritic cells (DCs), neutrophils and monocyte subsets (Ly6C$^{high}$ to Ly6C$^{low}$) of vehicle (corn oil) or TMX-treated wt or cKO mice with DSS colitis. Quantification of flow cytometric data (B) for frequency (C) and number of macrophages, DCs, Ly6$^{hi-mid\ or\ low}$ monocytes and neutrophils of vehicle (corn oil) or TMX-treated wt or cKO mice with DSS colitis.
(D) Colonic *Il10*, *Il1b*, *Il6* and *Tnf* mRNA expressions relative to *Actb* by qRT- of DSS-given vehicle (corn oil) or TMX-treated wt or cKO mice with DSS colitis.
